# Prolonged cell encapsulation and rapid filamented light biofabrication of muscle constructs in microgravity

**DOI:** 10.1101/2025.06.11.659059

**Authors:** Michael Winkelbauer, Jakub Janiak, Johannes Windisch, Hao Liu, Maria Bulatova, Max Von Witzleben, Hugo Oliveira, Sophie Dani, Richard Frank Richter, Nicolas L’Heureux, Ori Bar-Nur, Michael Gelinsky, Marcy Zenobi-Wong, Parth Chansoria

## Abstract

The prospects of fabricating human tissue grafts or models using cell-laden bioresins in space has garnered significant interest in recent years. While there has been tremendous progress in extrusion or light-based bioprinting in microgravity conditions, printing of aligned tissues, such as those featuring anisotropic organization of cells and extracellular matrices (e.g., muscle, tendon, cardiac, etc.), remains a challenge. Furthermore, current photoresin formulations do not allow long-term cell encapsulation and are difficult to perform in microgravity. In this study, we demonstrate a new gravity-independent filamented light (G-FLight) biofabrication system with in-built refrigeration and heating units, which can create viable muscle constructs within seconds. We developed new photoresin formulations based on gelatin methacrylate (GelMA) for encapsulation of primary cells (murine myoblasts) and storage in printing cuvettes for at least a week at 4°C or -80°C. The tissues printed in microgravity based on the new formulations exhibited higher cell viability, number of proliferating cells and after maturation higher numbers of myotubes and fusion index compared to control formulations (i.e., GelMA dissolved in phosphate buffered saline). The microgravity-printed tissues also featured similar myotube density and fusion index to those printed using the same resins on-ground. The G-Flight printing concept, together with the new resins enabling refrigeration or cryopreservation with encapsulated cells, offers a promising solution for biofabrication in space.

## 1. Introduction

The “space race” during the 1950-70s ushered in a bold and ambitious era of scientific innovation and technological breakthroughs, ultimately pushing the boundaries of what was thought possible in space travel.^1^ Despite a reduction crewed space missions thereafter, the spark for interplanetary exploration has persisted amongst the masses, and with the recent development of re-usable rockets and advanced computing systems which reduce costs and improve the efficiency of spacecrafts, such an exploration is becoming a tangible possibility.^2,3^ Sending humans into space, however, would require sustaining lives by understanding the pathophysiological changes to tissues in space (e.g., due the lack of gravity) and developing solutions to repair and regenerate human tissues damaged due to accidents, ageing and disease.^4–6^ Towards these applications, the field of biofabrication has an important role to play.^7,8^

Biofabrication tools have enabled both in-vitro and in-situ fabrication of engineered tissues and grafts which mimic the natural cellular and extracellular matrix environment of natural tissues.^8,9^ These tissues have been studied in space-like environments,^10^ including simulated microgravity (e.g., in random positioning machines),^11,12^ short-term microgravity (e.g., in parabolic flights),^13,14^ or prolonged microgravity (e.g., aboard the International Space Station (ISS)).^15–17^ While simulated microgravity approaches deploying continuous alteration of gravity vectors allows cost effectiveness and small form factor for prolonged experiments, current versions are more suitable for holding bioreactors and cell culture vessels.^8,12^ Testing a biofabrication approach in simulated microgravity would necessitate the development of a customized system for holding the bioprinter and will not recapitulate the actual printing environment and resin handling conditions. In contrast, parabolic aircraft maneuvers offer brief yet authentic microgravity, making them ideal for testing biofabrication systems before deploying them in cost-intensive space missions.

For a biofabrication approach suitable for space missions, the tissues could be either biofabricated directly under microgravity conditions, or biofabricated on-ground and then maturated in microgravity.^8,18^ Performing biofabrication in microgravity, as opposed to printing specific tissues on-ground, offers the opportunity for printing customized, patient-specific tissue structures with bespoke spatial distribution of cells and biomaterials catering to the complexity of the target tissues, which is highly relevant for future space exploration.^18^ Here, we hypothesize that a gravity-independent biofabrication approach can allow predictable processing of biomaterials similar to those fabricated on-ground. Such an approach should leverage rapid fabrication of tissues within seconds, which is typically achieved with light-based biofabrication techniques.^19,20^ Notably, techniques such as Xolographic and tomographic printing have been successfully used in microgravity conditions to fabricate complex shapes.^21,22^ Crucially, these studies demonstrated that microgravity environments facilitate the use of low-viscosity resins, which would otherwise be challenging to replicate on-ground, where gravity-induced sedimentation typically compromises the stability of printed structures..^18,20^ However, the demonstrations were limited to acellular resins, as cell encapsulation within resins introduces several material and logistical constraints,^8,10,18^ as we have catered to in our present work (more details below).

We have previously demonstrated (on-ground) filamented light (FLight) biofabrication as a rapid, biocompatible and scalable approach towards building mm-scale aligned tissue constructs mimicking the microarchitecture and matrix components of native muscle,^23^ nerve,^24^ and articular cartilage.^25^ In this work, we demonstrate a new gravity-independent FLight (G-FLight) printer which features a small and robust form-factor to fit into the aircraft and withstand vibrational and safety requirements (**Figure 1A**). As a necessary precursor to future space applications, we validated the G-FLight printer within parabolic flights which recreate several cycles of microgravity lasting 20-22s, which was the time window for printing the constructs.

**Figure 1.**
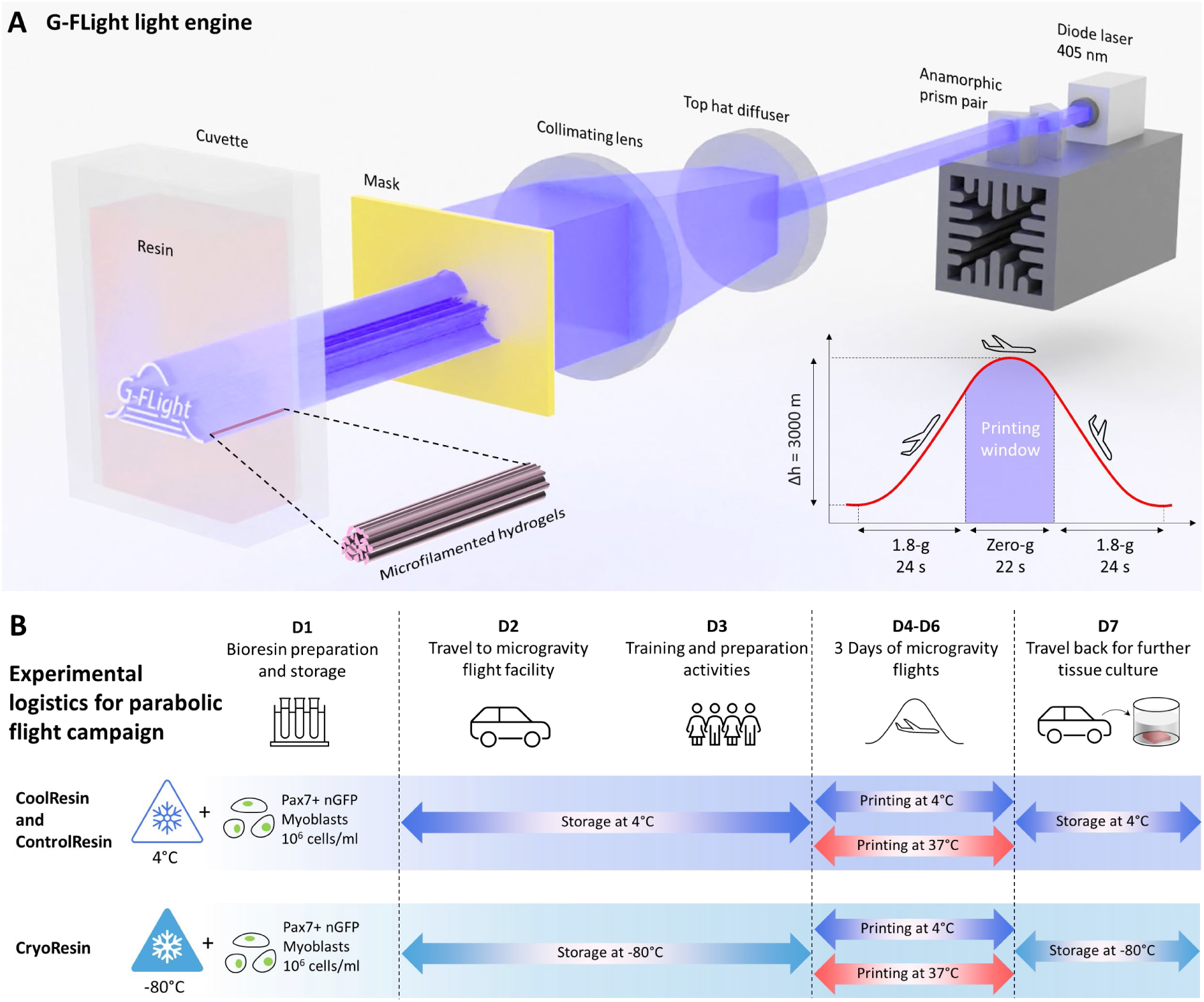
Optical components of G-FLight printer and resin formulations. **A.** Illustration of the components of the light engine in the G-FLight printer and the laser light path (also see **Video S1**). Here, the 405 nm laser beam was shaped into an elliptical format using the anamorphic prism pair and later converted into a square profile with a top-hat diffusor. A plano-convex lens then collimated the light, which was directed through custom photomasks onto a cuvette filled with the photoresin. This arrangement of components allowed for a robust and minimal form-factor while allowing rapid FLight printing. **B.** Logistics of the experiments which guided the resin storage and printing activities. Of note, the entire timeline of the activities spanned a week, and does not include the tissue culture period for 17 days which accompanied the parabolic flight campaign activities.

In addition to the considerations for the biofabrication system, the storage and handling of cell-laden bioresins also needed to be considered. Current bioresin formulations require specialized training for handling in microgravity.^26^ Here, we posited that the resin handling procedure can be simplified,^27^ if one could store the cell-laden resins in refrigeration (4°C) or cryopreservation (-80°C) conditions within sealed cuvettes (to ensure sterility), and, at the day of printing, simply thaw the vial (e.g., for the cryopreserved formulations) and print the tissues inside the cuvettes. Testing two different storage temperatures could allow the assessment of suitability for both short-term space missions (i.e., those lasting less than 2 weeks) which could suffice from a storage at 4°C or long-term missions (i.e., those lasting several months), which would necessitate cryopreservation (-80°C) conditions.

In this work, we developed new photoresin formulations – CoolResin and CryoResin – which allowed storage with encapsulated cells at 4°C or -80°C, respectively. These formulations were based on gelatin methacrylate (GelMA), as it is one of the most widely used biomaterials in light-based tissue engineering applications.^28–31^ The development of the new formulations was necessary to account for the experimental logistics (illustrated in **Figure 1B**), which involved several key steps: **1.** Cell encapsulation within the photoresins (i.e., to create the bioresins) followed by loading within the cuvettes (activities performed in the cell culture facility in Zurich, Switzerland), **2.** Establishing the appropriate storage conditions for the cuvettes (storage inside a portable refrigerator (4°C) or dry ice (-80°C)), **3.** Transportation to the parabolic flight facility in Bordeaux, France, followed by several days of training and preparation, and then parabolic flight days, during which the printing was carried out , **4.** Storage of the resins back at 4°C or -80°C after each day of printing, **5.** Driving back to the cell culture facility (Zurich, Switzerland), **6.** Retrieving the tissue constructs and culturing them to assess viability, cell differentiation and maturation characteristics.

Notably, as a control formulation (termed as ControlResin henceforth), we used GelMA dissolved in phosphate buffered saline (PBS), to be able to compare the cytoprotective effects and further tissue maturation for the CoolResin and CryoResin formulations. Furthermore, since the GelMA formulations exhibit a thermo-reversible gelation state at 4°C and a liquid state at 37°C, we investigated the impact of these two printing temperatures (also indicated in Figure 1B),^28,32^ to evaluate the resulting differences in constructs post-fabrication and maturation. We also printed constructs on-ground in tandem to the microgravity flights and compared them to the microgravity-printed constructs after maturation. We have discussed how these resin formulations can be deployed for use in future space applications for complex tissue biofabrication.

## 2. Results

### 2.1 Assembly of the G-Flight system for biofabrication under microgravity

The optomechanical assembly of the G-FLight system and additional auxiliary components were designed considering several engineering and logistical constraints. These included: **1.** Size constraint on the available rack space (30(l)×45(w)×30(h) cm) available in the aircraft performing parabolic flight maneuvers (see image of the rack in **Figure S1**), **2.** The need for an onboard resin refrigeration unit (for storage at 4°C during the parabolic flights) and heating unit, and **3.** The need to withstand the vibration during transportation and parabolic flight maneuvers. To address these constraints, we excluded sensitive optical components such as a digital micromirror device and complex beam-shaping optics (e.g., telescopic lenses), which we had used in our prior work,^23,33^ and instead developed a compact and robust light engine featuring 3D printed physical masks for light shaping (illustration in Figure 1A, animation of the light-path in Video S1). This system incorporated a 405 nm laser beam which was initially shaped from an elliptical profile (1×5 mm) into a rectangular profile (3×5 mm) using an anamorphic prism pair, and later into a square profile (10×10 mm) with a 20° divergence angle via a top-hat diffusor. A plano-convex lens was then positioned thereafter to collimate the light along a 30×30 mm profile, which was passed through custom-made photomasks to shape the light prior to projection onto a cuvette containing the photoresins (Figure 1A).

An illustration of the assembly of the G-FLight printer with all its auxiliary components (including the light engine inside an enclosure) is shown in **Figure 2A**, while the actual assembled printer and the key components are shown in **Figure 2B**. All the components were mounted on a 30×45 cm^2^ optical breadboard. The printer housed a refrigeration unit which stored the bioresins (i.e., cells encapsulated within the photoresins) between 4-6°C throughout the duration of the parabolic flight, and a heating unit for heating the resins to 37°C prior to printing. In the G-FLight printer, the print parameters (exposure duration and intensity) could be set using the graphical user interface (see **Video S2**). Prior to printing, appropriate masks could be retrieved from the mask holder and the photoresin-laden cuvettes retrieved from the heating or refrigeration unit and placed inside the printing enclosure. After printing, the uncrosslinked resin could be washed away (on-ground) and the printed constructs retrieved from the cuvettes when brought back to the cell culture facility (more details in the next section).

**Figure 2.**
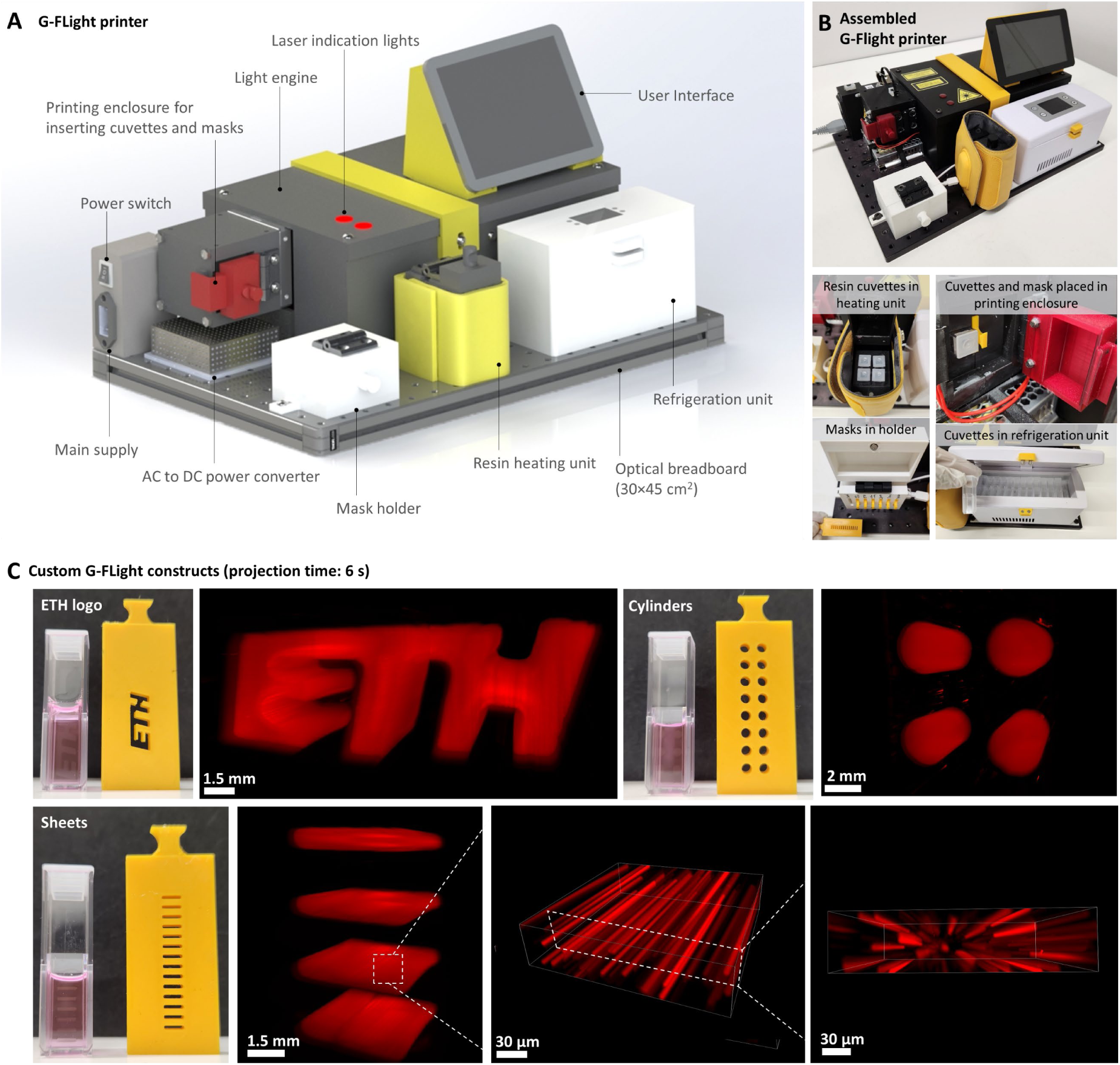
G-FLight printer and example prints of microfilamented constructs. **A.** Components of G-FLight printer contained within a volume of 30(l)×45(w)×30(h) cm. **B.** The assembled G- FLight printer and its key components used during the microgravity printing phase. **C.** Example prints of the ETH logo, cylindrical, and sheet-like constructs using rhodamine-labelled resins (ControlResin formulations containing GelMA at 5% w/v within PBS were used for these experiments; see methods for details). The inset images demonstrate the microfilaments within the G-FLight printed constructs. The images of the printed constructs are obtained through light sheet microscopy and images of the microfilaments within the constructs are obtained through confocal microscopy.

Representative prints using photoresins composed of 5% w/v GelMA dissolved in PBS are shown in **Figure 2C**. Based on the masks used, the light could be selectively shaped to control the cross-section of the constructs (examples of different masks and the corresponding prints are shown in Figure 2C). Upon reaching its gelation threshold, the illuminated photoresin locally crosslinked into hydrogels featuring micro-filaments – a characteristic of FLight-printed constructs – prevalent throughout the length of the hydrogels.

### 2.2 Development and characterization of photoresins for the encapsulation and storage of primary myoblasts under refrigeration and cryogenic conditions

Considering the experimental logistics (previously illustrated in Figure 1B), we developed the CoolResin and CryoResin resin formulations for storage with encapsulated cells at 4°C or - 80°C, respectively, and compared them to ControlResin formulations. The specific compositions of the resins are detailed in **Figure 3A**. All formulations contained 5% w/v GelMA in PBS with lithium phenyl-2,4,6-trimethylbenzoylphosphinate (LAP) as the photoinitiator at 0.1% w/v. For CoolResin, we supplemented the PBS solution with Hypothermosol^®^ FRS (termed as HTS henceforth) which is a commercially available defined hypothermic preservation medium.^34,35^ For CryoResin, we further supplemented (i.e., in addition to HTS) the PBS solution with trisaccharide melezitose hydrate and dimethyl sulfoxide (DMSO), which have been used previously in Cryo(bio)printing applications to prevent crystal formation in GelMA resins during freezing and improve cell survival.^36,37^ For the cells, we used satellite cell-derived Pax7-nGFP primary myoblasts,^33^ which express a GFP fluorescent reporter under a Pax7 promoter that is specific for muscle stem cells.^38,39^ These myoblast cells are known to activate upon injury *in vivo*, proliferate and differentiate, contributing to regeneration by forming new myotubes or replenishing the stem cell pool.^40,41^ These cells were especially suited to assessing the effects of the strenuous storage and transport conditions posed by the parabolic flight campaign. They also have been previously demonstrated to form biomimetic myotubes *in vitro* (upon induction of differentiation through serum withdrawal) exhibiting spontaneous contractions and sarcomere structures.^38,42^

**Figure 3.**
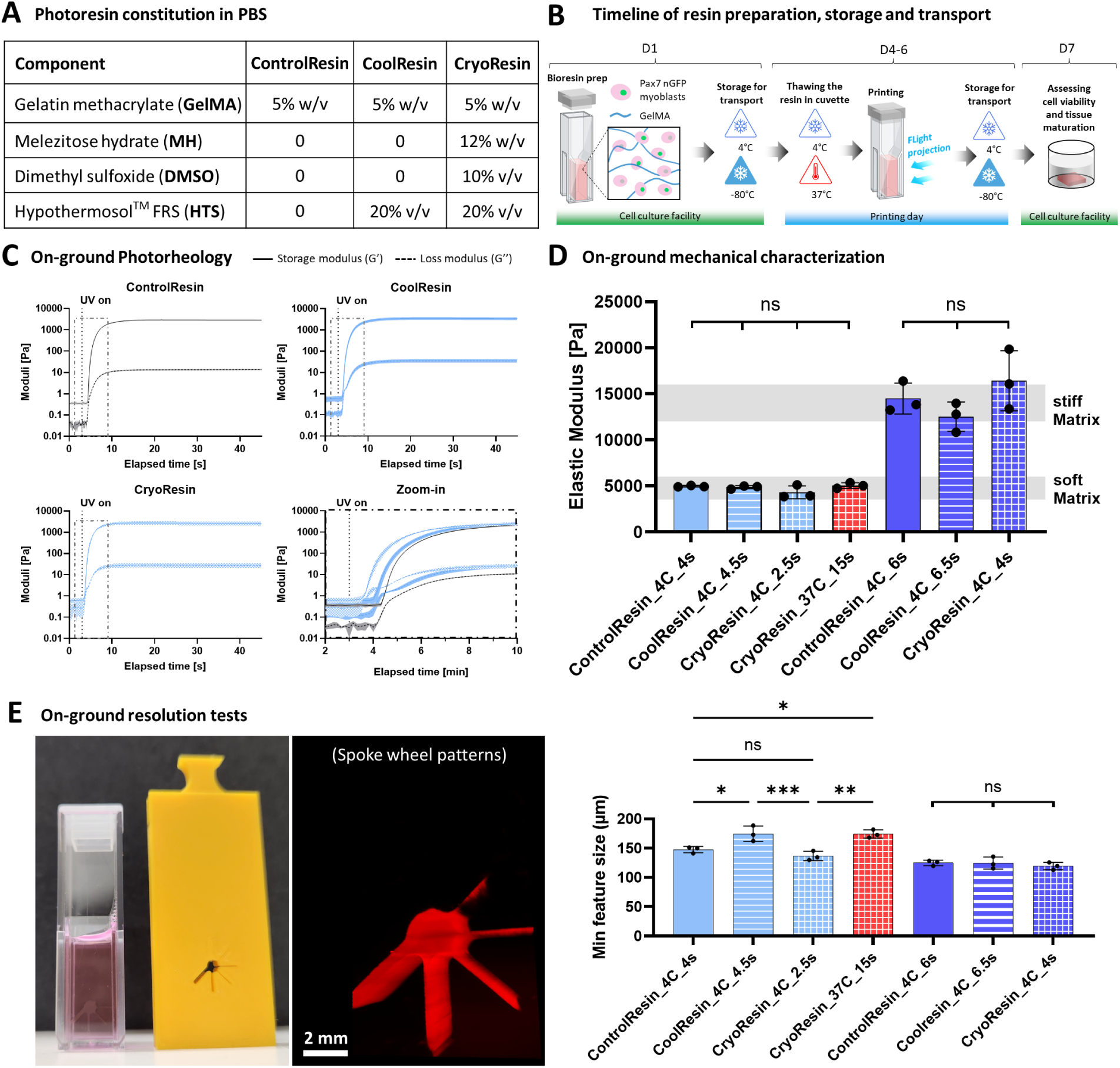
Experimental protocol and resin printing assessment. **A.** The components (dissolved in PBS) of CoolResin and CryoResin for cell-encapsulation and storage at 4°C or -80°C, respectively. The cellular resins were loaded within cuvettes and then stored for a week at 4°C or -80°C. In addition to these resins, we also included ControlResin formulations which consisted of GelMA mixed only in PBS. **B.** General timeline (spanning a total of 1 week) of resin storage, transport and printing. The resins (with cells) were transported from the cell culture facility in Zurich to the parabolic campaign facility in Bordeaux. On the day of printing, the resin formulations within the cuvettes were thawed and then printed using the G-Flight printer at 4°C or 37°C, and stored at 4°C after printing. The resins were then brought back to the ground and stored back at 4°C or -80°C until processed at the cell culture facility. **C.** Photorheology of the resin formulations, which demonstrated a rapid photocrosslinking under near UV (405 nm) light exposure. The zoom-in image shows the collective response of the resin formulations in the dotted region of the individual photorheology curves. **D.** Elastic modulus (under compression) of the printed constructs, where the exposure duration was changed to increase the crosslinking density in the matrix formulations. Through optimization of the light dose during polymerization, we selected specific formulations which were either soft (4-6 kPa) or stiff (12-16 kPa) for engineering the muscle constructs. **E.** Example prints using the resolution mask with the spoke wheel pattern and the analysis of the minimum feature size obtained using the different resin formulations. The labels 4C or 37C denote the printing temperatures 4°C and 37°C, respectively, while the timing (e.g., 4s, 6s, etc.) denotes the duration of FLight printing within the constructs, respectively. Data represented as mean ± SD (n=3), statistical significance was determined by one-way ANOVA and is denoted as follows: *refers to p<0.05, **refers to p<0.01, ***refers to p<0.001, ns refers to non-significant difference.

Pre-flight on-ground characterization of the crosslinking kinetics was performed after the storage of the resins for 1 week to account for the logistics of the parabolic campaign (**Figure 3B** highlights the timeline specific to the resins). Here, the CoolResin and ControlResin formulations were stored 4°C by simply transferring the sterile resin-filled cuvettes (with capped lids) in a refrigerator. For CryoResin formulations, the capped cuvettes with the resins were placed in a controlled-rate freezing unit (Mr. Frosty™) containing isopropyl alcohol (IPA), which was in-turn placed within a dry ice container, to ensure cooling at 1°C/min until -80°C. Note that the ControlResin and CoolResin formulations were not evaluated for storage at -80°C as the constructs demonstrated very poor fidelity and low cell viability (< 10%) after printing (pilot results not shown). The photorheological tests confirmed a rapid polymerization of GelMA for different formulations upon exposure to 405 nm light. To isolate the effects of resin additives from inherent physical crosslinking behavior of GelMA, these measurements were conducted at 25°C on non-gelled (liquid state) formulations. Comparative analysis of the crosslinking onset (**Figure 3C**) indicated faster crosslinking kinetics in the CryoResin formulations compared to the CoolResin and ControlResin, which could be attributed to inhibition of any photoinitiator degradation during cryogenic storage. Despite differences in polymerization onset, the final storage moduli after complete crosslinking were comparable, yielding values in the range 2.7-3.4 kPa.

Subsequently, we assessed the effect of photoexposure duration on hydrogel stiffness. Considering ranges used in muscle tissue engineering,^43^ we classified crosslinked hydrogels exhibiting an elastic modulus of 3-5 kPa as “soft” gels and 12–16 kPa as “stiff” gels (**Figure 3D**). Here, for the same light engine producing output at 34 mW/cm^2^, we screened a range of exposure times (see **Figure S2**). Notably, the resins being printed at 37°C took longer to polymerize and to achieve crosslinking ranges similar to those printed at 4°C, which can be attributable to the changes in molecular structure and reactive group availability in the GelMA formulations under cold or heated conditions.^32^ Notably, considering the printing window during the microgravity phase would be within 22 s, only the CryoResin formulations at 37°C (termed as CryoResin_37C henceforth) were suitable as they required a 15 s exposure duration. The resulting gels from CryoResin_37C exhibited stiffness values within the soft stiffness range. However, none of the other formulations (when heated to 37°C) reached the desired stiffness in either the soft or stiff regimes within the available 22-second printing window. Consequently, these formulations were excluded from subsequent cell-based experiments. From these tests, we obtained optimal exposure times as 4, 4.5, 2.5, and 15 s for ControlResin_4C, CoolResin_4C, CryoResin_4C, and CryoResin_37C, respectively, to cater to the soft stiffness regime (Figure 3C). For the stiff regime, prolonged exposure times of 6, 6.5, and 4 s were used for ControlResin_4C, CoolResin_4C, and CryoResin_4C, respectively (Figure 3D).The optimized light exposure durations were subsequently applied to evaluate the achievable printing resolution using the G-FLight system. For this, a spoke-wheel photomask (**Figure 3E**) was used, and the minimum achievable feature sizes were determined for the different formulations (**Figure S3**). In the soft stiffness regime, differences in the minimum achievable feature sizes were observed across formulations, with average resolutions of 147 ± 5.4 µm, 174.5 ± 13.22 µm, 136.6 ± 7.97 µm, and 174.1 ± 6.86 µm for ControlResin_4C, CoolResin_4C, CryoResin_4C, and CryoResin_37C, respectively (Figure 3E). Notably, constructs within the higher stiffness regime for each formulation exhibited a smaller minimum attainable feature size than those within the lower stiffness regime, while no significant differences in resolution were found amongst the formulations (minimum feature size ∼ 120 ± 5 µm). The inability to achieve smaller feature sizes with lower exposure duration is attributable to the spreading of light dose within the resin due to optical blurring and free radical diffusion, which necessitates the use of higher optical doses for smaller features. This is a known constraint with laser-based printing techniques such as tomographic printing, where the optical doses need to be digitally tuned to account for the different feature sizes in an object.^44^

Prior to characterizing the microarchitecture of the resin formulations and the characteristics of cells therein, we conducted pilot tests to screen the formulations for their ability to maintain shape fidelity during long-term culture. This was essential to be able to later study the muscle maturation characteristics. Accordingly, for these experiments, we encapsulated Pax7-nGFP primary myoblasts and cultured them for one week within both soft and stiff gels for the different formulations. Here, the the gels from the “soft” regime, except for the CryoResin_37C formulation, exhibited substantial change in their overall morphology, including warping and shrinking due to cell-matrix interactions (**Figure S4**). The maintenance of the fidelity for the CryoResin_37C formulation, despite being in the soft regime, could be attributed to the prolonged exposure duration (15 s) for these formulations, which can lead to non-specific crosslinking due to a combination of optical scattering and free radical diffusion.^45^ The ramifications of non-specific crosslinking in the CryoResin_37C formulation will be evident in the subsequent studies. Consequently, further microarchitectural characterization and long-term tissue maturation was limited to gels crosslinked to the stiff regime and the CryoResin_37C formulation, where no morphological alterations were observed during the pilot cell culture tests (Figure S4).

### 2.3 On-ground assessment of the microfilament characteristics and cell survival within the selected resin formulations

The G-FLight system used the FLight concept for the biofabrication of highly aligned microfilamented constructs. Importantly, the microarchitecture of FLight constructs is not only influenced by the fabrication technique (i.e., the light-engine), but also by the properties of photoresins used (e.g. refractive indices of the resin and concentration, etc.).^23^ For the selected resin formulations (**Figure 4A**) obtained after initial tests assessing construct morphology in culture (Figure S4), we performed confocal imaging to characterize the microfilament distribution within the constructs (**Figure 4B)**. Similar to the previous tests, the samples were stored for 1 week at 4°C (for CoolResin and ControlResin) or -80°C after controlled-rate freezing (for CryoResin).

**Figure 4.**
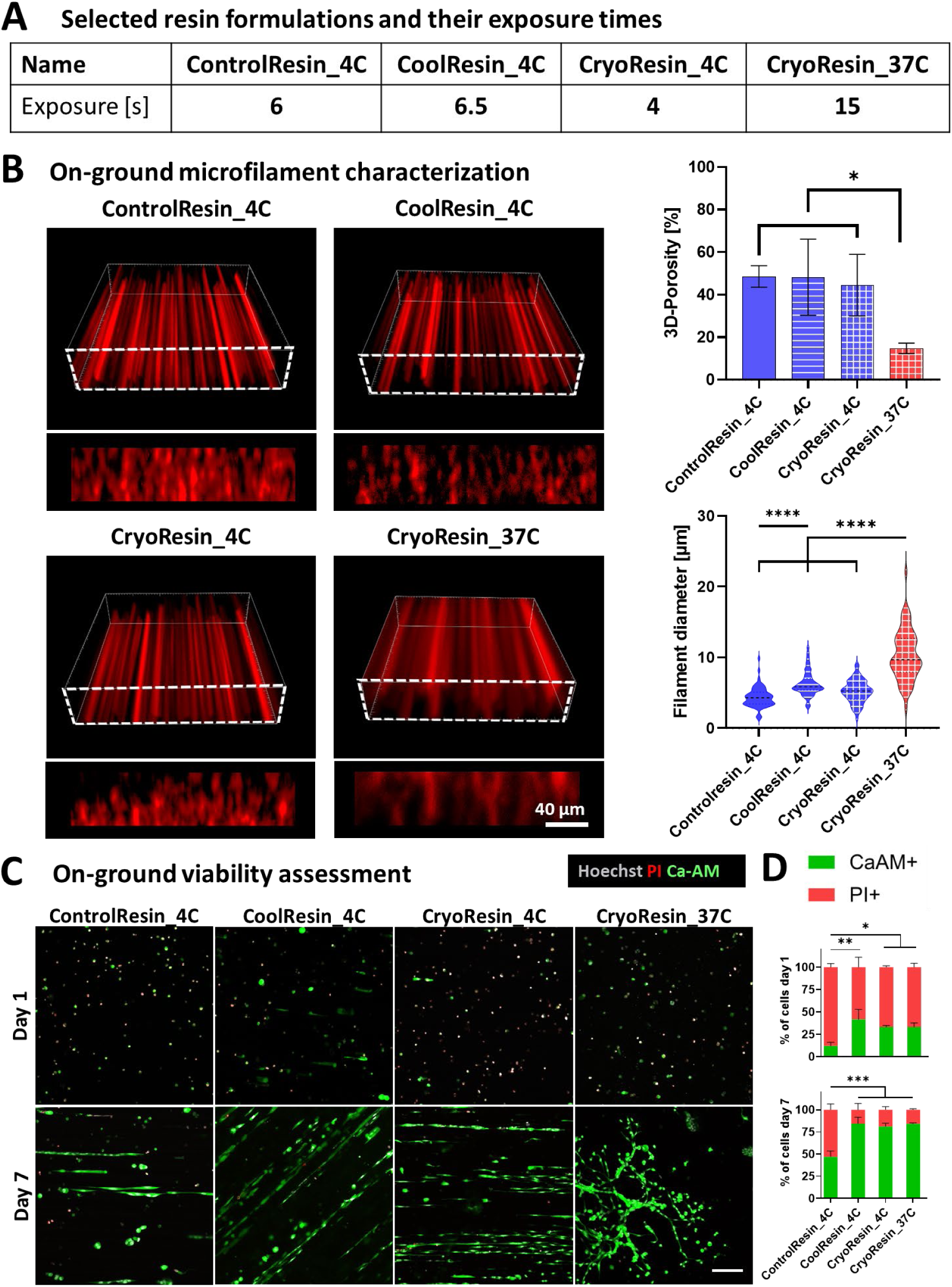
On ground pilot assessments of microfilament characteristics and cell survival within selected resin formulations. **A.** Naming convention (used henceforth) of selected resin formulations and their exposure duration. **B.** Microfilament distribution within the printed constructs using the selected resin formulations and the analysis of the microfilament size and porosity. **C.** Microscopic images of the cells stained with propidium iodide (PI) and Calcein AM. **D.** Analysis of the percentage of cells stained with Calcein AM and PI on days 1 and 7 of culture. Data represented as mean ± SD (n=3), statistical significance was determined by one-way ANOVA and is denoted as follows: *refers to p<0.05, **refers to p<0.01, **** refers to p<0.0001, ns refers to non-significant difference.

All resin formulations printed at 4°C (i.e.: in their solid state) displayed comparable 3D-porosity of approximately 50%, with similar distribution of microfilament diameters ranging from 1.5 to 12 µm. Specifically, the porosity (i.e., void space in the three-dimensional (3D) constructs) of constructs printed at 4°C were 48.6 ± 5.1%, 48.2 ± 17.8% and 44.5 ± 14.5 % and the microfilament diameter of 4.3 ± 1.4 µm, 6.2 ± 1.7 µm and 5.2 ± 1.6 µm for ControlResin_4C, CoolResin_4C, and CryoResin_4C, respectively (see naming conventions in Figure 4A). The Cryoresin_37C formulation (i.e.: in its liquid state) exhibited a significantly lower porosity of 14.8 ± 2.5% and a significantly broader distribution of microfilament diameters ranging from 2.9-to 22.1 µm with a diameter of 10.3 ± 3.5 µm. This low porosity and high variability in the microfilament diameters within CryoResin_37C formulation could be attributed to the prolonged exposure duration (15 s) and non-specific crosslinking as described previously.

After characterizing the microfilaments in the constructs, we encapsulated Pax7-nGFP primary myoblasts in the selected resin formulations at 10⁶ cells/mL. The bioresins were transferred into cuvettes and subjected to storage conditions specific to each resin: 4 °C for ControlResin_4C and CoolResin_4C, or –80 °C via a controlled-rate freezer for CryoResin_4C and CryoResin_37C. Considering the actual printing timeline (Figure 1B), after 4 days of storage, the cryopreserved cuvettes were rapidly thawed and transferred to 4 °C for two hours prior to light exposure and subsequent fabrication of cell sheets using the G-Flight printer. This step allowed emulating the conditions of the actual microgravity Flight campaign, where the cuvettes needed to be prepared before the flight and then placed in the onboard refrigerator within the G-Flight printer to be used during the parabolic flight maneuvers. The ControlResin_4C, CoolResin_4C and CryoResin_4C were printed at 4°C, while the CryoResin_37C were liquefied in the cuvettes using the onboard heating unit at 37 °C shortly before the printing. Following the printing procedure, all cuvettes were transferred to 4 °C for an additional 2 hours to emulate aircraft conditions. Finally, to simulate transportation of the cuvettes from the parabolic flight facility (in Bordeaux) to the laboratory (in Zurich), the cuvettes containing the bioresins were stored under their respective conditions: CryoResin-based samples were re-frozen to -80°C using controlled-rate freezing, and ControlResin and CoolResin-based samples were refrigerated at 4 °C for additional 3 days. Constructs were then rapidly thawed, washed extensively with PBS, and cultured in myoblast expansion medium for 7 days. Cell viability was assessed on days 1, 4, and 7 using CalceinAM (CaAM) and propidium iodide (PI) staining.

The selected images of the constructs from CaAM and PI staining are shown in **Figure 4C**, and corresponding data analysis is shown in **Figure 4D**.^[36]^ On Day 1, the highest proportions of viable (i.e., CaAM⁺) cells were observed in the tailored resin formulations including the CoolResin_4C (41.7 ± 11.2 %), CryoResin_4C (33.2 ± 1.8 %), CryoResin_37C (33.3 ± 4.3), which was significantly higher than the ControlResin_4C formulations (12.1 ± 4.1). The poor viability in the ControlResin_4C can be attributed to the absence of any HTS, which would otherwise act as a hypothermic preservation agent. Interestingly, on day 1, the formulations also contained some double-positive cells (CaAM⁺/PI⁺), indicating membrane-compromised cells (Figure 4C), which is consistent with literature.^46^ This could explain the less than ideal (we aimed for >80% viability on Day 1) viability observed on Day 1 for the CoolResin and CryoResin formulations. Nevertheless, by day 7, the proportion of viable cells had increased significantly, suggesting cellular recovery. Viability was assessed as 46.7 ± 6.6 %, 84.4 ± 7.1 %, 81.2 ± 3.7 %, and 84.2 ± 1.2 % for ControlResin_4C, CoolResin_4C, CryoResin_4C, and CryoResin_37C, respectively. The viability for the CoolResin and CryoResin formulations on Day 7 was significantly higher than the ControlResin formulations. Owing to the recovery of viable cells on Day 7 for the optimized resin formulations, the initial double-positive population likely represents cells in a stressed but viable state. We hypothesize that PI uptake occurred due to transient membrane permeability, rather than irreversible membrane damage, which is also typically observed during cryopreservation.^47^

### 2.4 Microgravity printing and assessment of myoblasts differentiation within the tissue constructs

Our on-ground analysis of the microarchitecture and cell survival allowed us to confirm the suitability of the resin formulations, in use with the G-FLight system, for subsequent printing in microgravity. For these experiments, the cells were encapsulated in the resins and loaded into cuvettes similar to the previous cell study. The cuvettes were then stored under their respective conditions at 4°C (portable refrigerator) for the ControlResin and CoolResin formulations, or at -80°C (using the controlled rate freezing procedure as described previously) for the CryoResin formulations.

On each flight day, the required cuvettes were retrieved and either directly transferred to the onboard refrigeration unit in the G-FLight printer (for ControlResin and CoolResin), or first rapidly thawed at 37 °C (for CryoResin) before transferring to the onboard refrigerator. Prior to each parabolic maneuver, the photomask (rectangular slits: 4(w)×1(h) mm) and the corresponding cuvette were installed in the printer. Each parabolic cycle (illustrated in **Figure 5A**) was initiated by a steep ascent, during which the aircraft experienced approximately 1.8 g for ∼24 seconds. Shortly before reaching the apex of the parabola, the crew signaled the onset of the microgravity window (∼22 seconds), during which the printing was performed while the aircraft was in freefall. After this zero-g phase, the aircraft entered a second hypergravity phase (∼24 seconds), completing one full parabolic maneuver. Between maneuvers, a 120–300 second interval in steady flight allowed for inspection of printed samples and preparation of the next print (i.e., loading a new cuvette and installing the corresponding photomask). A recording of the preparation, printing and inspection during one parabolic flight maneuver is shown in **Video S3**.

**Figure 5.**
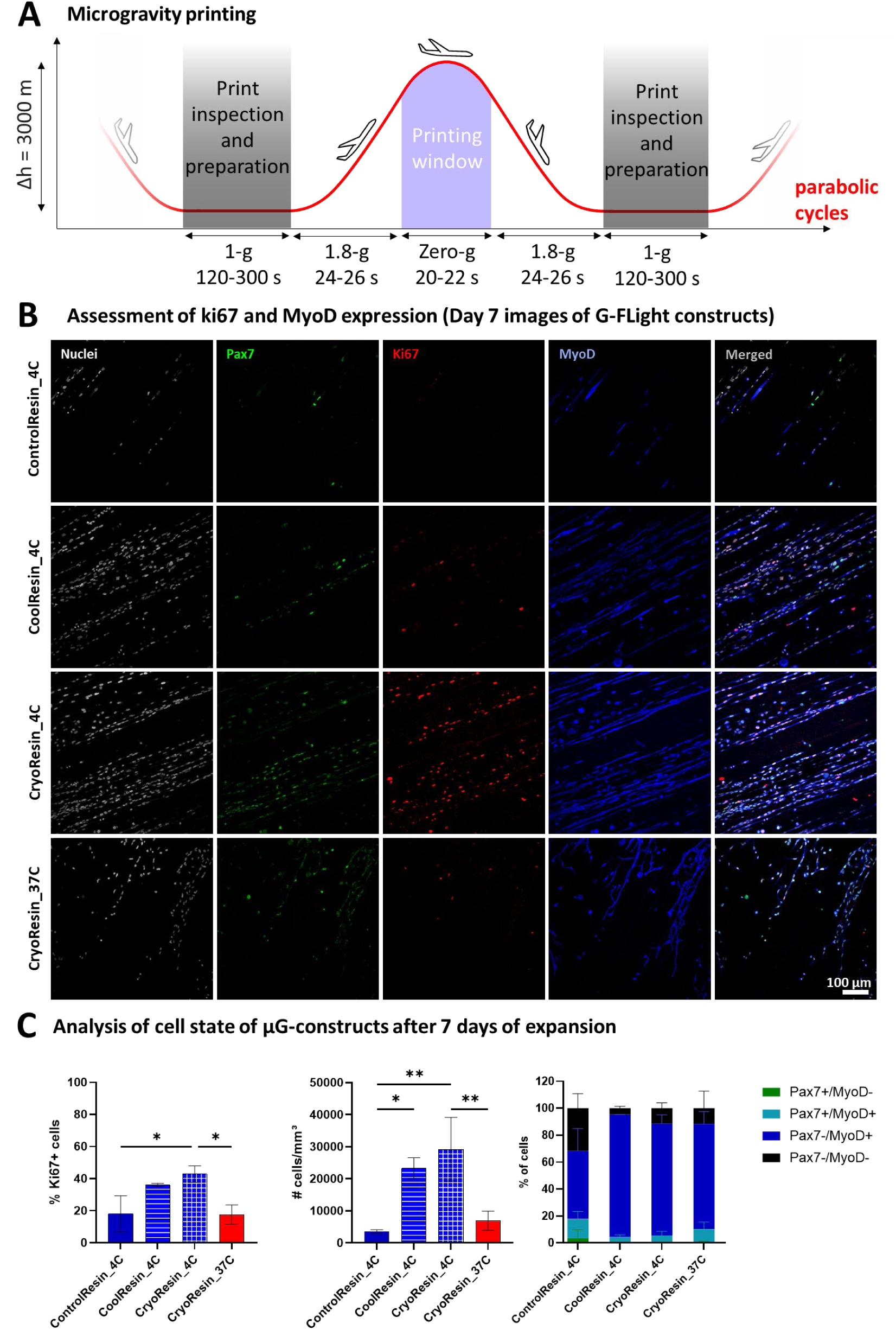
Microgravity printing activities and assessment of growth characteristics within the microgravity-printed constructs. **A.** The parabolic cycles (30 in total) consisted of several phases, where the cuvettes were retrieved from the refrigeration or heating unit, inspected and loaded into the printer during the 1 g (i.e., normal gravity) phase, and the prints executed during the 0 g (i.e., microgravity) phase. **C.** Microscopic images of the different resin formulations demonstrating Pax7^+^ (green), KI67^+^ (red) and MyoD positive (blue) cells (nuclei are grey). **C.** Analysis of the % of Ki67^+^ cells and accordingly, the #cells/mm^3^, and assessment of the myoblast differentiation state through the staining of Pax7^+^ and MyoD^+^ cells within the constructs after 7 days of culture. Data represented as mean ± SD (n=3), statistical significance was determined by one-way ANOVA and is denoted as follows: *refers to p<0.05, **refers to p<0.01.

Upon return to ground, the cuvettes were transferred back to their respective storage conditions (i.e. fridge or controlled-rate-freezer) until transportation to the cell culture laboratory at the end of the parabolic flight campaign. In the cell culture facility, the cuvettes with the printed constructs were thawed and washed thoroughly with PBS, followed by incubation in myoblast expansion medium. All samples were maintained in expansion medium for 7 days, after which the culture was switched to differentiation medium for 10 days to induce myoblast differentiation into myotubes. At the end of both the expansion and the differentiation period, constructs were fixed and processed using immunofluorescent staining.

At the end of the expansion period (i.e., day 7), we analyzed cells for Ki67 (i.e., proliferating cells), Pax7-nGFP (i.e., satellite cells with myogenic progenitor and self-renewal properties), and MyoD (i.e., cells differentiating towards myogenesis) expression (**Figure 5B).** Notably, uniaxial cell alignment along the microfilament direction was present in all the groups fabricated at 4°C. This alignment was missing in the CryoResin_37C formulation, which could be attributed to the non-specific crosslinking within these formulations which reduce the porosity and limit the cell guidance cues from the microfilaments. Quantification of proliferating cells (Ki67⁺) revealed that CoolResin_4C and CryoResin_4C supported the highest levels of proliferation, with 36.1 ± 1 % and 42.8 ± 5.2 % Ki67⁺ cells, respectively. In contrast, ControlResin_4C and CryoResin_37C showed markedly lower proliferation, with only 18 ± 11.3% and 17.6 ± 6 % Ki67⁺ cells, respectively. The significantly lower amounts of cells in the ControlResin_4C group can be attributed to the absence of components for hypothermic preservation or cryopreservation, which caused reduced cell viability and overall cell count. The corresponding lower amounts of Ki67⁺ cells in the CryoResin_37C can be attributed to the low porosity of the matrix, which can limit cell proliferation and impede alignment. Owing to the high number of proliferating (Ki67⁺) cells, the total observed cell densities (including both Ki67^+^ and Ki67^-^ cells) were significantly higher in the CoolResin_4C and CryoResin_4C constructs, reaching 23.3 ± 3.2×10^3^ and 29.2 ± 9.9×10^3^ cells/mm³, respectively, compared to 3.5 ± 5.9×10^3^ and 6.9 ± 3×10^3^ cells/mm³ in the ControlResin_4C and CryoResin_37C formulations, respectively (**Figure 5C**). Importantly, these findings with the G-Flight printed constructs were consistent with our on-ground printing experiments, where the amount of Ki67⁺ cells and the observed cell densities at Day 7 were higher in the CoolResin_4C and CryoResin_4C, while being significantly lower in the ControlResin_4C and CryoResin_37C formulations (**Figure S5**).

To further characterize the cell state, we assessed MyoD expression and used Pax7-nGFP fluorescence signal to identify muscle stem/progenitor cells (Pax7^+^/MyoD^-^ and Pax7^+^/MyoD^+^) as well as committed cells (Pax7^-^/MyoD^+^). Although no statistically significant differences in MyoD⁺ or Pax7⁺ expression were detected between groups, the trends were consistent with on-ground experiments (Figure S5). Notably, from the on-ground experiments (Figure S5B), we observed elevated Pax7 expression during the early culture period, with approximately 40% of cells being Pax7⁺ or Pax7⁺MyoD⁺, indicating activation and self-renewal of the encapsulated myoblasts. By day 7, Pax7 expression had decreased, likely due to continued proliferation and increased cell densities prior to the switching to the differentiation medium. Nevertheless, the increased number of proliferating cells and the cell density within the CoolResin_4C and CryoResin_4C demonstrated the promising potential of these formulations in creating contractile muscle constructs, as explored in the subsequent studies.

### 2.5 Assessment of myotube characteristics in the microgravity-printed constructs after maturation

Differentiation of the constructs was initiated on day 7 by switching from expansion medium to differentiation medium. Myogenic maturation was assessed after an additional 10 days in culture via immunofluorescent staining for myosin heavy chain (MyHC) and filamentous actin (F-actin). Staining confirmed the fusion of myoblasts into multinucleated myotubes across all groups (**Figure 6A**). The myotube diameter for ControlResin_4C, CoolResin_4C, CryoResin_4C, and CryoResin_37C was 9.9 ± 3.3 µm, 12.6 ± 2.6 µm (significantly higher than that for ControlResin_4C), 11.9 ± 3.3 µm, and 11 ± 2.9 µm, respectively. Myotube density was found highest in CoolResin_4C and CryoResin_4C at 115 ± 0.2×10^3^ and 1.2 ± 0.2×10^3^ myotubes/mm³, respectively, while significantly lower densities were observed in ControlResin_4C (0.5 ± 0.1×10^3^ myotubes/mm³) and CryoResin_37C (0.7 ± 0.1×10^3^ myotubes/mm³). These differences can be explained by the significantly higher initial cell numbers prior to differentiation in the former two groups (Figure 5C). Using the MyHC signal, we further calculated the fusion index (i.e., the number of nuclei within myotubes divided by the total number of nuclei) and found similar trends, with the highest fusion indices in the CoolResin_4C and CryoResin_4C constructs (**Figure 6B**).

**Figure 6.**
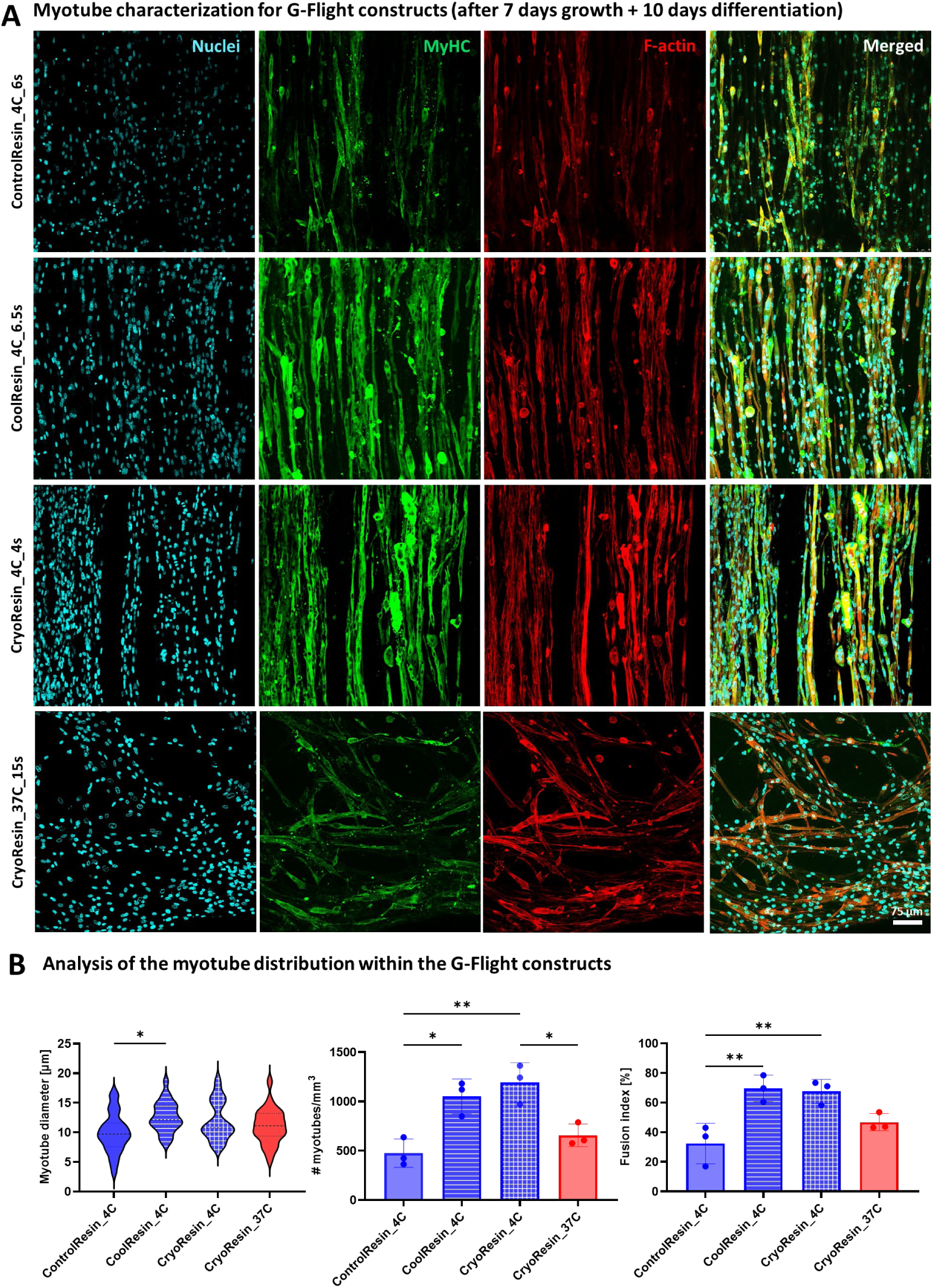
Myotube characteristics in the microgravity-printed constructs. **A.** All constructs demonstrated myotube formation after 17 days in culture (1 week of growth + 10 days of differentiation). Notably, the alignment of the myotubes in all formulations except for Cryoresin_37C formulation was uniaxial, i.e., in the direction of the microfilaments. **B.** The constructs featured an average myotube diameter between 10-15 µm, with the Controlresin_4C formulations demonstrating the lowest average diameter amongst the formulations. The number (#) of myotubes/mm^3^ and the fusion index were the highest within the CoolResin_4C and CryoResin_4C formulations) and porosity of the constructs (compared to the Cryoresin_37C formulations). Data represented as mean ± SD (n=3), statistical significance was determined by one-way ANOVA and is denoted as follows: *refers to p<0.05, **refers to p<0.01.

Collectively, these results indicate that CoolResin_4C and CryoResin_4C supported the most robust myogenic differentiation and maturation. Importantly, these two groups also demonstrated similar myotube characteristics to those printed on-ground (**Figure S6**). These groups were therefore analysed further using electrical response tests and sarcomere staining.

### 2.6 Assessment of sarcomere structures and contractile response of selected microgravity-printed constructs after maturation

Based on the previous findings, we selected the best-performing conditions (CoolResin_4C and CryoResin_4C) for further analysis of functional muscle maturation within the microgravity-printed constructs. Spontaneous myotube contractions were observed as early as day 12 of culture (i.e., day 5 of differentiation) in these two formulations. Constructs were stained for sarcomeric α-actinin (SAA) to visualize the basic contractile units of skeletal muscle, the sarcomeres (**Figure 7A**). Morphological analysis revealed well-aligned, striated myotubes with regular sarcomere spacing of ∼ 2.7 µm (**Figure 7B**), which lies within the range typically found in mouse muscle.^48^ To assess functional contractility, we evaluated the response of the constructs to external electrical stimulation. Constructs were placed in fresh medium and oriented with platinum electrodes positioned aligned to the longitudinal axis of myotubes. A square wave stimulus was applied at 1 V/mm with 40 ms pulse duration at stimulation frequencies of 1 Hz and 5 Hz (**Figure 7C**). Myotube contractions were recorded via live imaging, and the displacement amplitude was quantified (see **Video S4** and **Video S5** showing contractions at 1 Hz and 5Hz, respectively). Notably, constructs responded synchronously to the electrical stimulation. At 1 Hz, the average displacement amplitude was ∼4 µm, while at 5 Hz, the amplitude increased to ∼7 µm, consistent with the onset of tetanic contractions. These results confirm that Pax7-nGFP myoblasts, when encapsulated in the optimized resin formulations and stored for up to one week and printed in microgravity, retain their capacity to form structurally mature and functionally contractile muscle tissue constructs.

**Figure 7.**
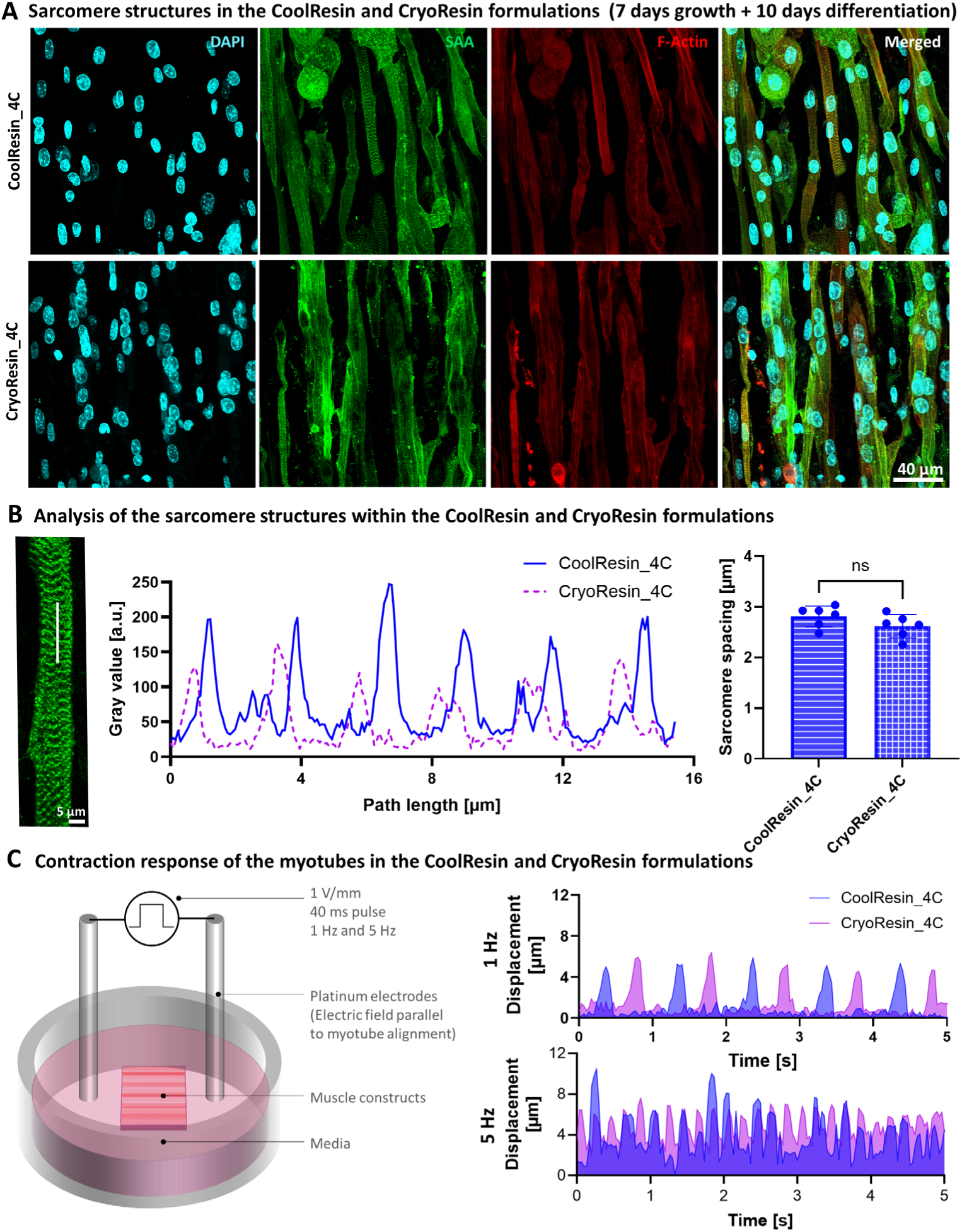
Characterization of the sarcomere structure and contractile response within the microgravity-printed muscle constructs. **A.** The constructs made from Coolresin_4C and CryoResin_4C formulations stain positive for sarcomeric alpha actin staining and demonstrate regularly interspaced sarcomere structures. **B.** Analysis of the sarcomere structures (gray values are plotted along a selected line (white color) over the sarcomere images) from both formulations reveals similar sarcomere spacing of ∼2.8 µm within the constructs. **C.** The printed constructs demonstrate spontaneous contractility and synchronization to external electrical stimulus (apparatus) providing pulse electrical pulses of 1 V/mm amplitude at 1Hz and 5Hz (also see Videos S4 and S5). Note that the initiation of the excitation was shifted by approximately half the frequency to be able to observe construct response for each formulation more clearly. Here, the muscle fibers within the constructs made using both CoolResin_4C and Cryoresin_4C formulations demonstrate similar contraction amplitude and synchronization to external stimulus.

## 3. Discussion

Biofabrication of engineered tissues and grafts can help sustain long-term space missions and provide insights on disease mechanisms in space.^7,8,10^ In this work, we focused on FLight as gravity-independent approach for the rapid biofabrication of aligned muscle tissues. We have used the term “gravity-independent” in context with the similarity of the microgravity-printed tissues to those fabricated on-ground (Figure S6). This is largely enabled due to the rapid (i.e., within seconds) fabrication using the G-FLight system and the GelMA-based resin formulations where the cells were confined in a viscous hydrogel material (especially within the formulations printed at 4°C). The characteristics of the maturated muscle tissues (e.g., myotube diameter, fusion index and sarcomere spacing) printed using the G-FLight system are comparable to existing approaches using casting,^49–51^ micropatterning,^52,53^ and the native mouse muscle.^54^ These attributes support the potential use of G-FLight system for the fabrication of muscle constructs in space, with notable use in disease modeling,^55^ or as regenerative grafts.^56^ For instance, one of the unfavourable attributes of space travel is rapid atrophy of skeletal muscle.^57,58^ Here, the printed muscle constructs can be used as platform to screen for means to maintain or regenerate muscle mass or prevent its loss in microgravity (small molecules, exercise biomimetics etc.),^59^ as a potential next exploratory step. Other growing research avenues including biohybrid muscle actuators,^60^ or cultivated meat,^61^ can also be potentially explored with this method.

While this study focused on Pax7⁺ nGFP primary mouse myoblasts, which are known for their high regenerative capacity and resilience to environmental stress,^62,63^ it remains to be determined whether these resin formulations and the G-FLight system are broadly compatible with other, potentially more sensitive, human cells (e.g., induced pluripotent stem cell-derived cells).^64,65^ The favourable results observed here indicate that the optimized resin formulations and the printing approach support survival, proliferation, and differentiation even after storage and processing. The printing approach and materials could therefore also be deployed towards other tissues such as cartilage,^25,66^ tendon,^67^ neurons,^24^ or cardiovascular,^68^ which have been previously printed on-ground using FLight biofabrication.

In this work, the use of physical masks to shape the light patterns allowed miniaturization of the light engine in the G-FLight system, to be able to conform to the dimensions of the airplane rack. However, physical masks limit the flexibility of the shapes that can be fabricated; for instance, hollow tubular shapes are not possible with the current system since the inner circle of the tube will be a free-floating structure within the 3D printed mask. If a larger size of the printing system may be permissible in the future, the use of digital micromirror device (DMD) for light shaping could obviate this limitation.^23,25^ Furthermore, FLight printing, in its current form, is limited to 2.5D prints, where the cross-section is constant throughout the fabricated structures.^20,69^ Here, other deep vat printing approaches such as tomographic printing,^70,71^ or light sheet-based printing (e.g., Xolography),^72–74^ can be deployed using the same bioresins which we have shown in this work, to produce complex 3D constructs. Recently, the projection of multi-wavelength light patterns has allowed selective photocrosslinking or photodegradation within multimaterial bioinks.^69,75,76^ These strategies enable the on-demand fabrication of complex multi-cellular tissues, reinforcing the rationale for conducting tissue biofabrication in space rather than transporting pre-fabricated tissues on space missions. Nevertheless, being able to culture pre-fabricated tissues in space is highly relevant for disease modeling. For such studies, the tissues could be fabricated on-ground, followed by perfusion with media containing MH, HTS and DMSO to enable long-term cryopreservation and transport to space,^8^ where the preservation media could be removed and the tissues cultured further in bioreactors.

The CoolResin and CryoResin formulations we demonstrated in this work allowed storage of cell-laden resins for at least a week under refrigeration or cryopreservation conditions. In these formulations, the use of HTS allowed preservation of the cells under hypothermic conditions, while the use of MH and DMSO prevented crystal formation in the cryopreserved formulations. These material constituents can possibly also be transferred to other formulations, such as those based on collagen,^42,77^ hyaluronic acid,^78^ silk,^79,80^ decellularized matrix,^81^ or fibrinogen,^82^ etc., which have been widely used in tissue engineering and regenerative medicine. These formulations have been used as non-modified (e.g., using native tyrosine crosslinking) or functionalized (e.g., methacrylated, or norbornene and thiol-functionalized) with FLight or volumetric printing,^25,42,79,83^ and can expand the range of tissues which can be created. There is also ongoing research in the field of cryopreservation, including the development of alternative cryoprotectants and optimized freezing protocols.^84,85^ In this context, the CryoResin formulation presents a promising platform for long-term storage (several months to years). The resins could be implemented with broadly used cryopreservation techniques, potentially enabling wider application across different cell types and off-the-shelf biofabrication workflows. Beyond viability, assessing changes in gene expression and downstream effects after long-term storage will offer valuable insight into how cells retain their functional potential for tissue-specific applications. Finally, the transfer of cryopreserved (–80 °C) constructs into liquid nitrogen (–196 °C), followed by successful retrieval and activation, would further demonstrate the feasibility of using such formulations in long-duration space missions.

Future space exploration applications may also necessitate the use of multiple biofabrication approaches in conjunction with one another. This would leverage the advantages of each approach and overcome its limitations through a hybridization scheme.^86–88^ The resin formulations used in this work could be adapted for use in conjunction with other technologies, such as extrusion printing or two-photon polymerization, broadening their applicability across diverse fabrication platforms.^88,89^ Importantly, when using light-based biofabrication systems in space, as a standalone or hybrid approach, one must consider the steps for washing away the uncrosslinked resin and culturing the tissues under perfusion. In future, we will investigate the design of customized print containers which could allow easy coupling to perfusion systems for removal of the uncrosslinked resin followed by media flow to facilitate tissue maturation.^18^ Notably, such perfusion-based culture systems can also be tested under simulated microgravity, i.e., tissues can be fabricated in parabolic flights followed by maturation under perfusion in simulated microgravity (e.g., in RPM machines).^12^ However, there are differences in the cellular adhesion, mechanosensitive gene expression and mitochondrial function in the cells when comparing simulated microgravity to real microgravity such as in space flights.^90,91^ Therefore, biofabrication and long-term culture within real microgravity, such as aboard the ISS,^15,16^ will also be investigated in our future work.

## 4. Conclusion

In this work, we have presented G-FLight printing as an effective tool for the rapid gravity-independent fabrication of aligned tissues, focusing on muscle tissue as an application. The printer featured a robust and compact form-factor capable of printing aligned microfilamented tissue constructs within seconds. We optimized specific resin formulations which allowed storage together with encapsulated cells under refrigeration or cryopreservation conditions for at least one week. The printed constructs from optimized resin formulations exhibited increased cell survival (>80% viability) compared to the controls (< 50% viability) after a week in culture. In the maturated tissue constructs, we observed higher myotube density and fusion index compared to controls. Finally, we also observed sarcomere structures in the tissue constructs made from the optimized formulations, with spontaneous myotube contractility and synchronization to external stimulus. By demonstrating gravity-independent FLight printing together with cell-laden resin formulations which allow long-term storage, this study can pave the way for new light-based approaches for tissue biofabrication in space.

## 5. Methods

### 5.1 Assembly of the G-FLight printer

The printer components were mounted onto an optical aluminum breadboard (30×45 cm^2^, MB3045/M) containing M6 taps spaced-out by 25 mm. These taps were used to mount all the printer components including the cover to allow a robust assembly. The refrigeration unit for the storage of the resins at 4°C during the parabolic maneuvers was a commercially available insulin storage unit box utilizing a Peltier cooling system, and the heating unit consisted of a custom 3D printed Polyethylene terephthalate glycol (PETG) enclosure with a 3D printed cap and an aluminum jacket wrapped around for even heat distribution. The heat was generated by wrapping a mini flask warmer (Thermup Go, Lionel Care) which was set to 41°C, which allowed maintenance of 37°C inside the heating unit to liquefy the resins prior to printing (only for the formulations which were printed in a heated state). The other printer components, including the mask holder, masks, printing enclosure and the cover of the printer were also custom 3D printed using PETG and mounted onto the optical breadboard.

As for the optical and electrical components, a supplementary figure has been provided detailing the components and their connections (see **Figure S7**). The laser module (LDM-405, LaserTack GmbH) was purchased as a collimated source with anamorphic pairs mounted at the output to generate a square beam profile (4×4 mm^2^) at 300 mW. A top hat diffusor (ED1-S20-MD, Thorlabs) was placed along the laser light path to homogenize the beam (i.e., remove the gaussian intensity distribution) at the output of the laser. A plano convex lens (f = 75 mm, Thorlabs) was then used to collimate the light beam which was subsequently shaped using the masks inserted into the printing chamber. The optical components were covered by an aluminium enclosure to prevent light leakage as per the safety norms dictated by NoveSpace. Additional laser indication lights were connected to the laser controller to turn ON when the laser was activated. In addition, a reed switch was mounted onto the printing closure (where the masks and cuvettes were inserted) and was interlocked with the laser controller. A magnet was attached to the lid of the printing enclosure, which, upon closing, activated the laser. The laser controller received its analog signals through a Raspberry Pi 4.0 which in-turn received user input for the light intensity and the print duration through a graphical user interface displayed on a touch screen. All the components were powered by a single DC power supply unit (12 V, 50 W, LRS 50-12, Mean Well) which was connected to the mains supply. The components running on 12V (Laser module and refrigeration unit) were directly connected to the power supply (with appropriate fuses), while the components requiring 5V supply (Raspberry Pi and heating unit) were interfaced with a 12V-5V converter (LM2596, Purecrea, Bastelgarage) between the component and the power supply.

### 5.2 Photoresin constitution

Gelatin methacryloyl (GelMA) with a high degree of functionalization (DoF) was synthesized following our established protocol. Briefly, Type A gelatin (G2500, Sigma) was dissolved at 10% (w/v) in 0.25 M carbonate–bicarbonate buffer (pH 9) at 50 °C under constant stirring. Methacrylic anhydride (MAA, 276685-500, Sigma) was added in five equal portions over the course of 2 hours (one addition every 30 minutes), maintaining vigorous stirring and constant pH (9.0) throughout the reaction. A total MAA-to-gelatin ratio of 0.4 mL per gram of gelatin was used. One hour after the final addition, the reaction was terminated by 2-fold dilution with deionized water and pH adjustment to 7.4. The solution was centrifuged to remove unreacted MAA and then dialyzed (12–14 kDa MWCO, 3110, Merck) against Milli-Q water for 5 days with frequent water changes. The purified solution was sterile filtered (0.22 µm) and lyophilized. GelMA was stored at –20 °C until use.

To quantify the DoF, both unmodified gelatin and GelMA were dissolved at 1% (w/v) in deuterium oxide and analyzed by ¹H NMR (Bruker Ultrashield 400 MHz, 64 scans), see **Figure S8**. Spectra were normalized to the aromatic proton peaks of phenylalanine (7.1–7.5 ppm), which remain unaltered during functionalization. The ε-methylene protons of lysine (2.95–3.05 ppm) were integrated and compared between gelatin and GelMA to calculate the DoF using equation **(1)** and found to be 99%.

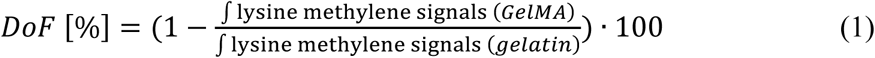

The ControlResin formulations were created by simply dissolving lyophilized GelMA at 5% w/v in PBS and allowing dissolution of the resin for 30-45 min at 50°C. The resin was allowed to cool down to 37°C followed by adding LAP (Lithium phenyl-2,4,6-trimethylbenzoylphosphinate) as the photoinitiator. For constituting the CoolResin or CryoResin formulations, dissolution solutions were first prepared by mixing HTS (for CoolResin) or a combination of HTS, MH and DMSO (for CryoResin) at the desired concentrations (**Figure 1B**) first in PBS and later adding GelMA at 5% w/v and LAP at 0.1% w/v (same dissolution procedure as the ControlResin formulations). The resins were freshly prepared the day before cell encapsulation or any in vitro evaluation and were stored at 4°C until used. Prior to usage, the resins were liquefied at 37°C for 5 min followed by further processing (i.e., encapsulation of cells or addition into cuvettes).

### 5.3 Cell culture, bioresin constitution and tissue maturation

The Pax7-nGFP mouse primary myoblasts and related culture methods were kindly supplied by Prof. Ori Bar-Nur. To grow and expand myoblasts in 2D conditions, the culture flasks were pre-coated with a coating solution which was made by mixing 1 ml of Matrigel (Corning, 356237) with 24 ml of low-glucose DMEM (Thermo Fisher Scientific, 31885023) and 1% v/v Penicillin-Streptomycin (Thermo Fisher Scientific, 15140122). After adding the coating solution, the culture flasks were incubated in the fridge for 7 minutes, coating solution was collected and flasks incubated at 37 °C for a minimum of 1 hour before use. A myoblast expansion medium comprised of Dulbecco’s Modified Eagle Medium (DMEM, Thermo Fisher Scientific, 41966029) and F-10 Medium (Thermo Fisher Scientific, 22390025) in equal parts, which was further supplemented with 10% v/v horse serum (ThermoFisher, 16050122), 20% v/v fetal bovine serum (Gibco, 10270106), 1% v/v Penicillin-Streptomycin (ThermoFisher Scientific, 15140122), and 10 ng/mL recombinant FGF2 (Bio-Techne, 233-FB-500). The differentiation medium comprised of 1% v/v Penicillin-Streptomycin (ThermoFisher Scientific, 15140122), 1% v/v Insulin-Transferrin-Selenium (Gibco, 41400-045), and 2% v/v horse serum (Thermo Fisher Scientific, 16050-122) added to DMEM (Thermo Fisher Scientific, 31966-021).

Upon reaching 80% confluency, the cells were harvested using Trypsin-EDTA 0.25% (ThermoFisher, 25200056) treatment, centrifuged at 500 g for 4 min, and suspended in the different resin formulations at 10^6^ cells/ml. After FLight printing (on-ground or after bringing back from the parabolic campaign), the hydrogel constructs were first cultured in expansion medium for 7 days (37 °C with 5% CO_2_), with the medium refreshed every 2 days. The tissue constructs were then incubated in differentiation medium for 10 days, where the media was replaced every 2 days, and once spontaneous muscle contractions were detected, the medium was changed daily.

### 5.4 Photoresponse characterization of the resin formulations

Photorheology experiments were conducted as per our previous study,^92^ using a rheometer (MCR 302e, Anton Paar) fitted with a 20 mm serrated plate geometry and a glass base. A UV lamp (Omnicure Series 1000, Lumen Dynamics) was used along with narrow 405 nm bandpass filters (Thorlabs). To prevent sample drying, a moist tissue paper was placed in the chamber during testing. Oscillatory measurements were performed in triplicates at 37 °C, using 152 µL of photoresin, with a 2% shear rate, 1 Hz frequency, a 200 µm gap, a 10-second acquisition interval, and an intensity of 2 mW/cm^2^.

### 5.5 Mechanical testing of the crosslinked constructs

Unconfined compression testing was conducted using a Texture Analyzer (TA.XTplus, Stable Micro Systems) fitted with a 0.5 kg load cell. After positioning the sample between two compression plates, a pre-load of 0.3 grams was applied to ensure complete contact with the samples. Following a relaxation period, the samples were compressed to 15% strain at a rate of 0.01 mm/s. The loading and unloading curves were recorded, and the compressive modulus was determined by performing a linear fit on the stress-strain curve as per our previous work.^93^

### 5.6 Light sheet imaging of the printed constructs

Light sheet imaging of the construct-laden cuvettes was performed as per the methods described in our previous work.^94^ Briefly, acryloxyethyl thiocarbamoyl Rhodamine B (Rhod-Acr) stock in DMSO (at 10 mg/mL) was added to the resin formulation at 1 µl/mL to enable fluorescence imaging. For facilitating imaging, the constructs were printed in 4 mm path length glass cuvettes (Thorlabs), which allowed the constructs to stick to the cuvettes.^94^ The constructs were washed 4 times with warm PBS at 37°C to remove any uncrosslinked resin. A light sheet microscope with axial scanning capability (MesoSPIM, V4) was employed to capture images of fluorescently labeled samples. The cuvettes with the constructs were placed on the MesoSPIM microscope stage and imaging was conducted using a macro-zoom system (Olympus MVX-10) and a 2x air objective lens (Olympus) with adjustable zoom functionality. For each imaging session, voltage adjustments were made using an electrically tunable lens (ETL). The step size for imaging ranged between 10 and 50 µm.

### 5.7 Microfilament imaging and size assessments

The rhodamine-labeled resins (method for fluorescent labeling are described in the previous subsection) were imaged using a confocal microscope. To quantify the 3D porosity of the hydrogels, confocal image stacks were converted into Imaris-compatible file formats using the Imaris File Converter software (Oxford Instruments, Ver. 10.2.0), and subsequently imported into Imaris (Oxford Instruments, Ver. 10.2.0) for surface reconstruction. Filament surfaces were rendered using the software’s standard surface detail settings and default intensity thresholds, unless stated otherwise. This analysis was performed for three different hydrogel matrices, and the filament volume obtained from surface reconstructions was used to compute 3D porosity.

For microfilament diameter characterization, the same confocal image stacks were processed using Fiji (ImageJ, Ver 1.54m). Z-projections of the top, middle, and bottom regions of each stack were generated, and filament diameters were determined by measuring the full width at half maximum (FWHM) of the resulting intensity profile.

### 5.8 Live/dead assessments of the muscle constructs

For viability assessments on Days 1 and 7, the cells in the constructs were stained with Hoechst 33342 (Invitrogen, 1:1000), propidium iodide (PI, Fluka, 1:500), and CalceinAM (Invitrogen, 1:2000). For assessment, a confocal laser scanning microscope (Fluoview 3000, Olympus) was utilized to capture 100 µm z-stacks with a 3 µm step interval (n=3). Image stacks were converted to Imaris-compatible format, imported into Imaris, and analyzed using the spot detection function to identify and count total nuclei and PI⁺ nuclei. The percentage of PI⁺ was calculated as the number of PI⁺ cells divided by the total number of nuclei. The percentage of Calcein-AM⁺ cells was calculated as: 100 – (% PI⁺ cells).

### 5.9 Immunohistochemistry and analysis of the muscle constructs

The constructs were fixed using 4% v/v paraformaldehyde for 30 minutes at room temperature, followed by washing with PBS. Prior to staining, the cells in constructs were permeabilized by treating the constructs with 0.2% v/v Triton X-100 in PBS for 15 minutes, followed by three washes in PBS. Blocking was performed with 5% w/v bovine serum albumin (BSA) in PBS for 1 hour. The constructs were subsequently incubated with either a primary anti-myosin heavy chain antibody (MF-20, DSHB, anti-mouse, 1:20 in PBS + 1% v/v BSA) or an anti-alpha-actinin (sarcomeric) antibody (A7811, Sigma, anti-mouse, 1:500 in PBS + 1% v/v BSA), or anti-ki67 antibody (550609, BD BioSciences, anti-mouse 1:200 in PBS + 1% v/v BSA) together with anti-MyoD antibody (PA5-23078, Invitrogen, anti-rabbit, 1:100 in PBS + 1% v/v BSA) for 24 hours under constant agitation. After three additional 15 min PBS washes, the constructs were incubated with appropriate secondary antibodies: Goat anti-mouse (AlexaFluor488, Invitrogen, 1:500 in PBS + 1% v/v BSA) and/or Goal anti-rabbit (AlexaFluor647, A-21244, Invitrogen, 1:500 in PBS + 1% v/v BSA). Hoechst 33342 (H3570, Invitrogen, 1:1000 in PBS + 1% v/v BSA), and phalloidin-tetramethylrhodamine B isothiocyanate (P1951, Invitrogen, 1:1000 in PBS + 1% v/v BSA) at 4°C for 2 hours. Finally, the samples were washed three times in PBS before being imaged using confocal microscopy (Fluoview 3000, Olympus).

### 5.10 Assessment of contractile response of the muscle constructs

For stimulation, the matured tissue constructs for the CoolResin and CryoResin formulations were transferred into 12 well plates filled with 1 ml of differentiation media. Separately, on the lid of the 12 well plates, 1 mm diameter holes were drilled and 1 mm diameter platinum electrodes inserted (17.5 mm spaced-out). The electrodes received electrical stimulus through an external power source which was interfaced to an Arduino via a TIP120 transistor. The voltage on the power source was set to achieve a 1V/mm voltage gradient parallel to the fiber orientation. The Arduino code was set to stimulate the muscle constructs using 4 ms pulses at 1 or 5 Hz. The setup was then placed atop a brightfield microscope (M5000, EVOS) and the video recordings performed at 60 fps using an external camera. The recordings were taken of both the spontaneous and electrically-responsive contractions of the muscle constructs. The plots for the displacement of the myotubes were determined using a custom python script analyzing the video frames based on our existing study.^42^

### 5.11 Statistical analysis

Statistical analysis was carried out in GraphPad Prism (v. 10.2.3) using One-way ANOVA followed by Tukey’s post-hoc analysis to compare groups. A significance level of 0.05 was applied. Statistical significance is indicated as follows: *p* < 0.05 (*), *p* < 0.01 (), *p* < 0.005 (*), and *p* < 0.001 (); “ns” denotes no significant difference.

## Supporting information

Laser light patch in the light engine

Demo of the G-Flight printer

Using the G-FLight printer during a parabolic flight (faces removed)

Stimulation of muscle constructs at 1Hz

Stimulation of muscle constructs at 5Hz

## Supporting information

A supporting information file and supplemental videos have been provided by the authors.

## Acknowledgements

M.W. and P.C. acknowledge support by the Swiss National Science Foundation (Ambizione grant PZ00P2_216356). M.G. and P.C. acknowledge support from The German Space Agency at DLR (grant number: 50WB2332) for funding the 43^rd^ parabolic flight campaign (PFC 43) in Bordeaux (France) organized and executed by NoveSpace. The authors also acknowledge the support from NoveSpace personnel for their engineering, technical and safety support for successfully executing the experiments in this work. The authors further acknowledge the MESOSPIM initiative at University of Zurich for the light sheet microscopy facility, and ScopeM at ETH Zurich for confocal microscopy facility.

## Data availability statement

All data is available in the ETH online repository with DOI: 10.3929/ethz-b-000739779.

