## Supplementary material for "Prolonged cell encapsulation and rapid filamented light biofabrication of muscle constructs in microgravity": Figures S1 to S8

This file contains **Supplementary Figures S1 to S8**

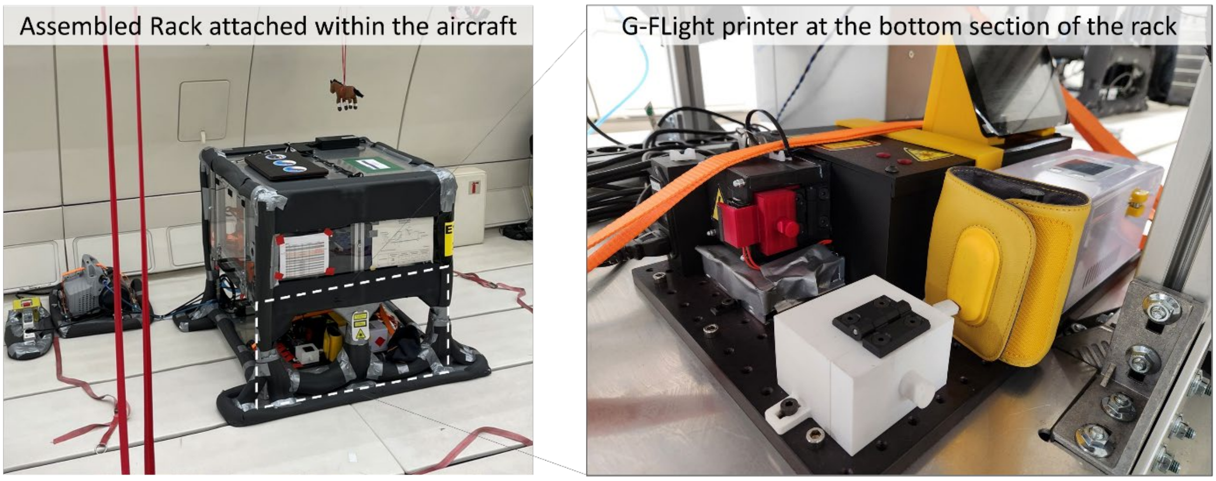


**Figure S1.** Photograph of the rack installed within the aircraft (left) and photograph of the printer attached within the rack (right).

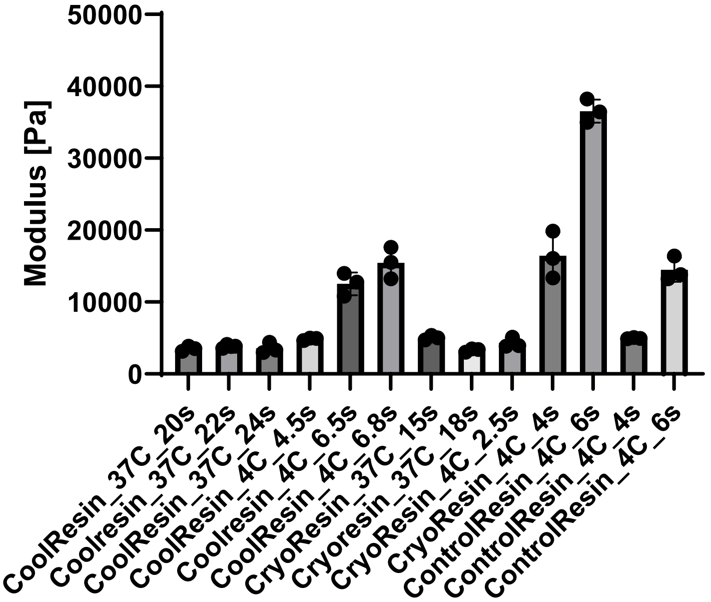


**Figure S2.** Compressive modulus of the printed formulations under different exposure duration using the same light engine (34 mW/cm$^2$).

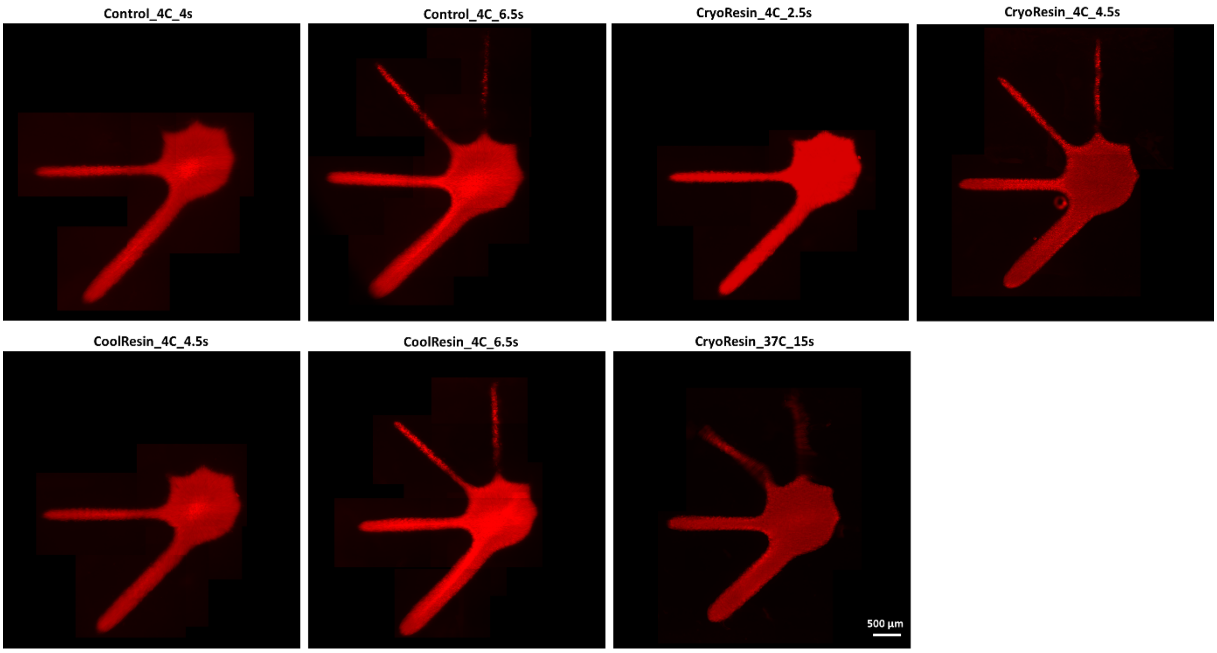


**Figure S3.** Resolution images of the selected resin formulations. Of note, the longer exposure durations allow emergence of thinner features within the constructs.

**A Construct morphology after a week in culture**

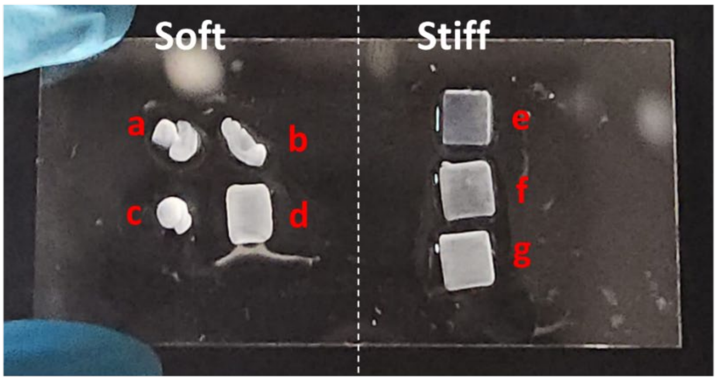


| ID | Group |
|---|---|
| a | ControlResin_4C_4s |
| b | CoolResin_4C_4.5s |
| c | CryoResin_4C_2.5s |
| d | CryoResin_37C_15s |
| e | ControlResin_4C_6s |
| f | CoolResin_4C_6.5s |
| g | CryoResin_4C_4s |

**B Cell morphology in the soft constructs which demonstrated matrix warping (other constructs are shown in the main manuscript)**

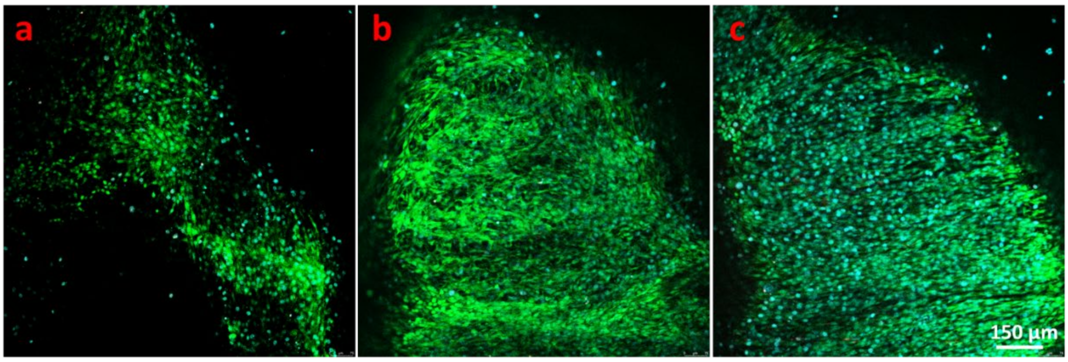


**Figure S4. A.** The printed constructs (image on the left) and their compositions (table on the right) after a week in culture. **B.** The morphology of the soft constructs (The resin compositions for the groups a, b and c can be found in the table in A), which demonstrates significant warping within the ControlResin and CoolResin formulations.

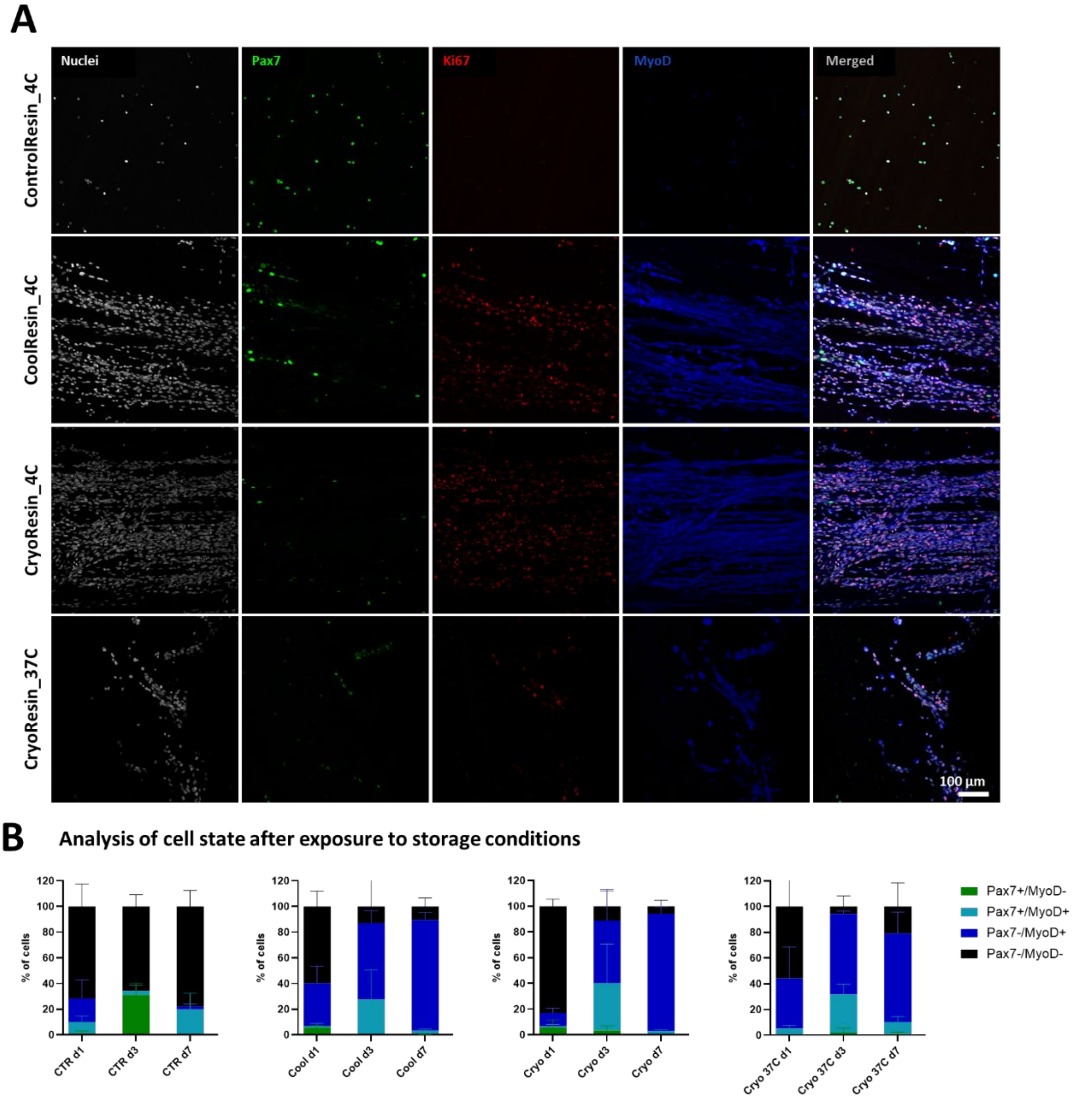


**Figure S5. A.** Staining for nuclei (grey), Pax7+ (green), ki67+ (red) and MyoD+ (blue) signals within the constructs after a week in culture. **B.** Respective analysis of the overlap of the Pax-nGFP and MyoD signals within the constructs over a week in culture. The abbreviations CTR, Cool, Cryo and Cryo37C refer to Controlresin_4C, Coolresin_4C, Cryoresin_4C and Cryoresin_37C.

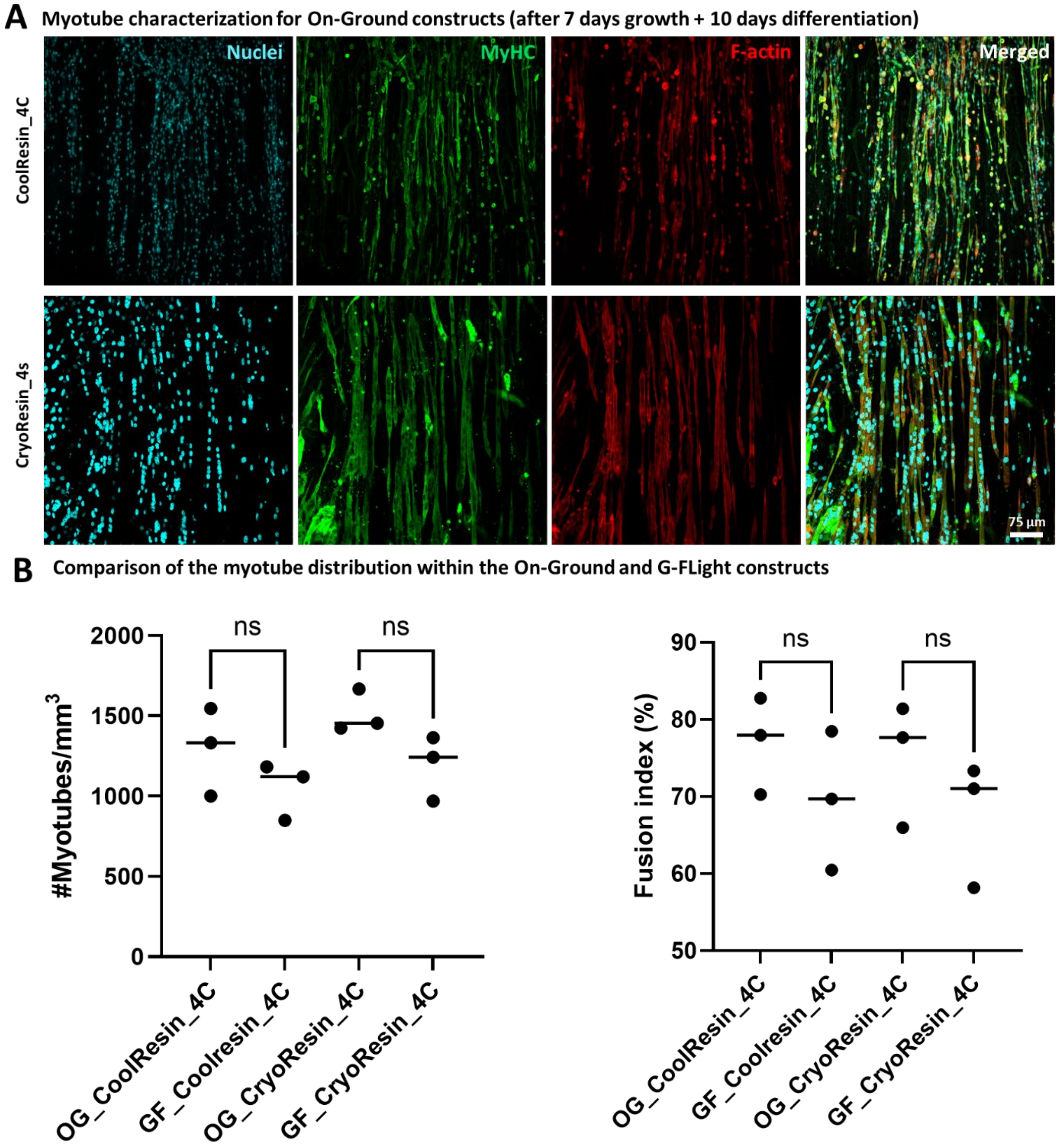


**Figure S6. A**. Staining for myosin heavy chain within the constructs printed On-Ground after maturation (7 days expansion + 10 days differentiation). **B.** Comparison of G-Flight (abbreviation GF) and On-Ground (abbreviation OG) printed constructs using the CoolResin_4C and Cryoresin_4C formulations shows no difference in the myotube density and fusion index within the matured constructs for each formulation.

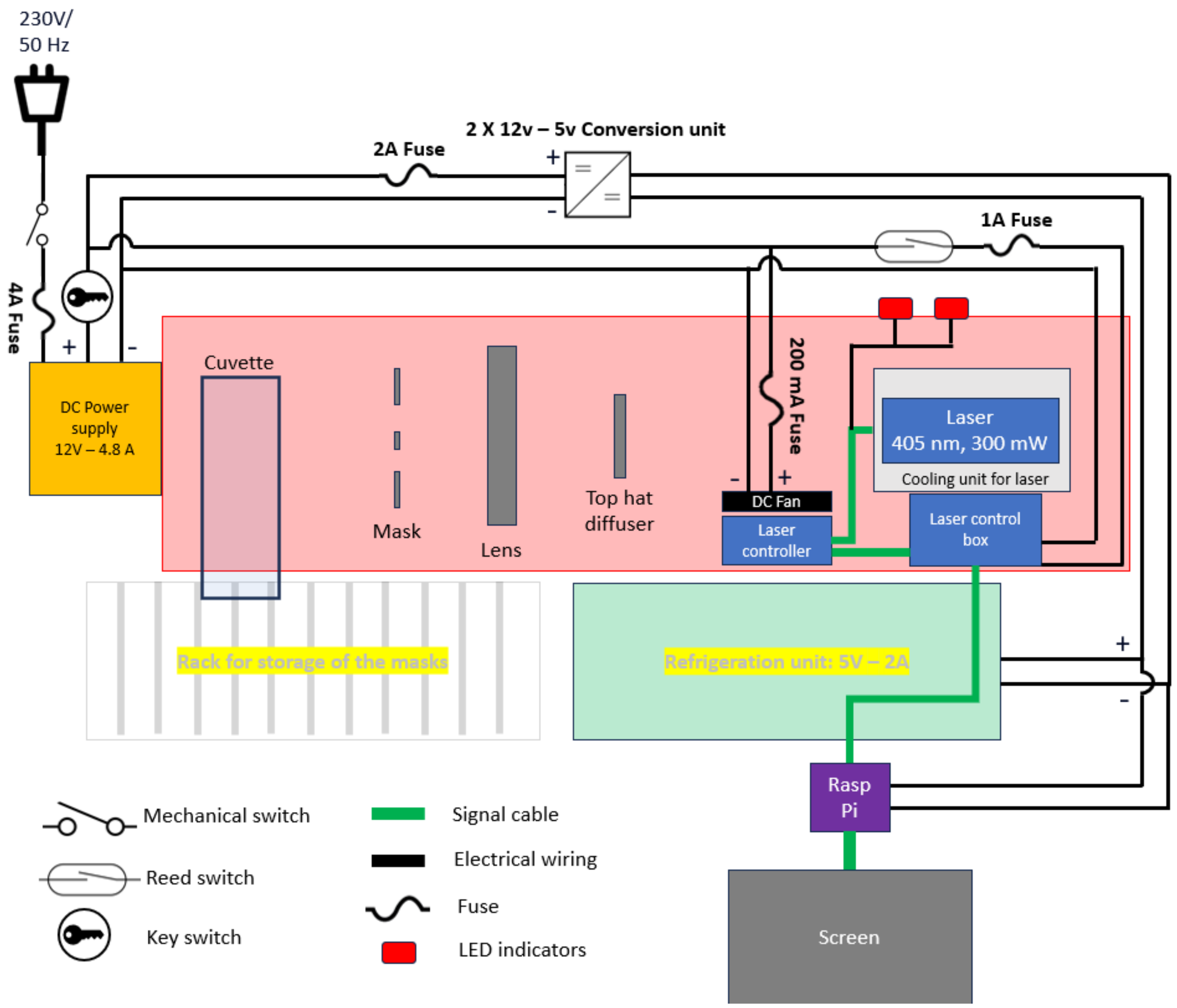


**Figure S7.** Component layout of the G-FLight system and corresponding electronic circuitry.

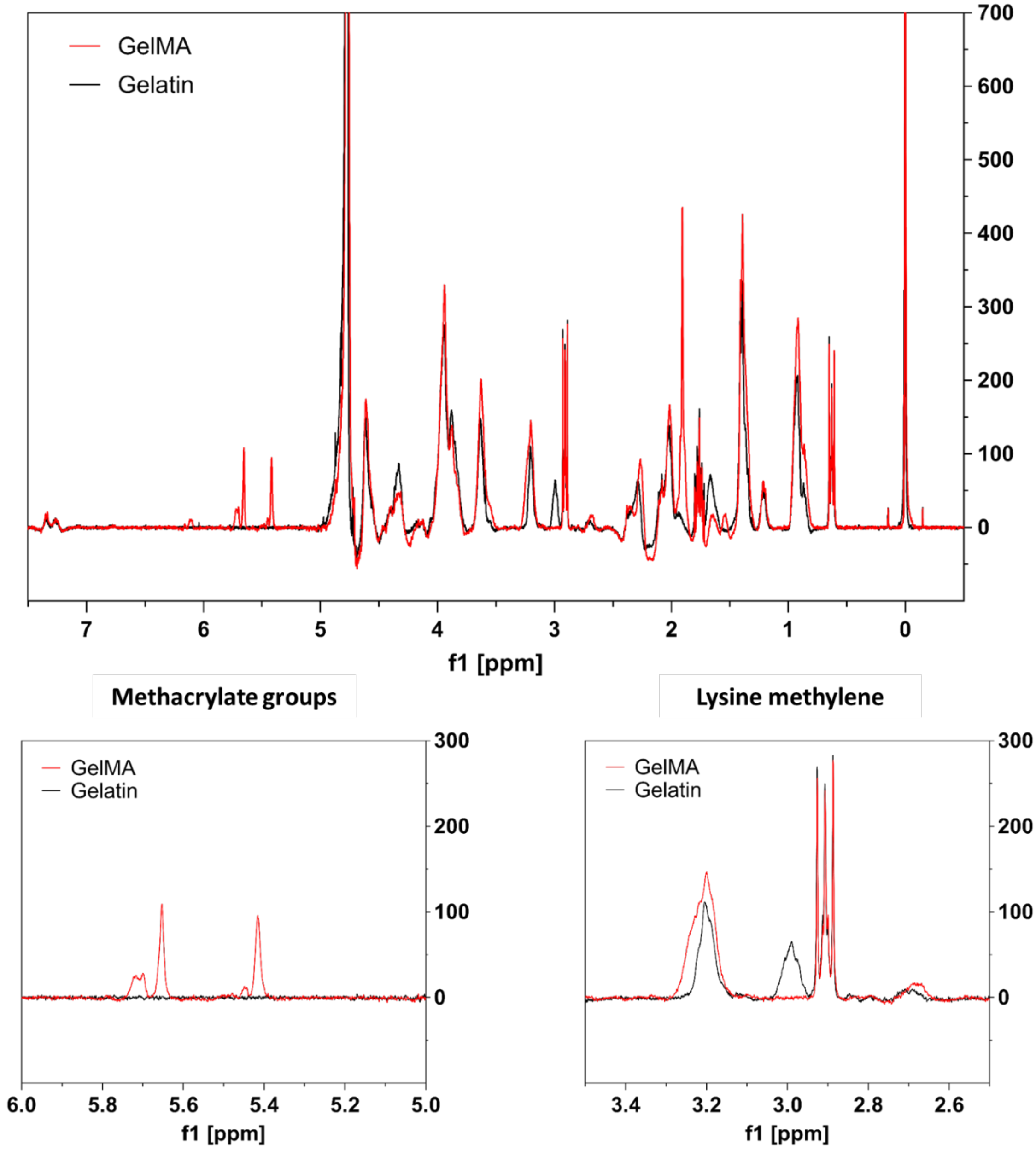


**Figure S8.** $^{1}$H NMR spectra of the GelMA formulation compared to pure gelatin.
